# A plant virus causes symptoms through the deployment of a host-mimicking protein domain to attract the insect vector

**DOI:** 10.1101/2022.12.16.520777

**Authors:** Man Gao, Emmanuel Aguilar, Borja Garnelo Gómez, Laura Medina-Puche, Pengfei Fan, Irene Ontiveros, Shaojun Pan, Huang Tan, Edda von Roepenack-Lahaye, Na Chen, Xiao-Wei Wang, David C Baulcombe, Eduardo R Bejarano, Juan Antonio Díaz-Pendón, Masahiko Furutani, Miyo Terao Morita, Rosa Lozano-Durán

**Author notes:** These authors contributed equally to this work (Man Gao, Emmanuel Aguilar).

## Abstract

Viruses are obligate intracellular parasites with limited proteomes that heavily rely on the cell molecular machinery for their multiplication and spread. Plant viruses frequently cause symptoms through interference with host developmental programs. Despite the agricultural relevance of symptom development in virus-infected crops, the molecular mechanisms underlying these viral effects remain elusive. Here, we show that the symptoms triggered by tomato yellow leaf curl virus (TYLCV) depend on the physical interaction between the host-mimicking domain of a virus-encoded protein, C4, and a plant-specific family of RCC1-like domain-containing (RLD) proteins. C4 outcompetes endogenous interactors of RLDs, disrupting RLD function in the regulation of endomembrane trafficking and polar auxin transport, ultimately leading to the developmental alterations recognized as symptoms of the viral infection. Importantly, symptoms do not have a detectable effect on the performance of the virus in the plant host, but they serve as attractants for the viral insect vector, the whitefly *Bemisia tabaci*, hence promoting pathogen spread. Our work uncovers the molecular underpinnings of the viral manipulation that leads to symptom development in the TYLCV-tomato pathosystem, and suggests that symptoms have evolved as a strategy to promote viral transmission by the insect vector. Given that most plant viruses are insect-transmitted, the principles described here might have broad applicability to crop-virus interactions.

## MAIN TEXT

Viruses fully rely on the cells they infect for subsistence and continuity. A successful viral infection and the concomitant manipulation of the host molecular machinery by the invading virus often lead to alterations in host physiology and/or development, which are recognized as symptoms. In plants, viral symptoms can include strong developmental changes that often dramatically reduce the productivity of infected crops, e.g. stunting, organ malformation, and chlorosis (Osterbaan & Fuchs, 2019). However, despite the obvious agricultural impact of virus-caused symptoms, the molecular underpinnings causing their appearance are not understood.

Different models have been proposed to explain the development of symptoms during infection by plant viruses. The competitive disease model advocates for the virus-induced diversion of limited resources towards viral processes as the basis for these developmental alterations (Culver & Padmanabhan, 2007). The frequent lack of correlation between symptom severity and viral load, however, favours the alternative interaction disease model, which proposes that specific interactions between virus and host disrupt the latter’s normal physiology and development, resulting in symptoms (Culver & Padmanabhan, 2007). Along these lines, viral symptoms have also been proposed to derive from the ability of the pathogen to interfere with RNA silencing (Wang *et al*., 2012). Nevertheless, whether symptoms confer a competitive advantage to the virus or, on the contrary, are merely side-effects of virulence activities is still an open question.

The family *Geminiviridae* (geminiviruses) comprises insect-transmitted plant viruses with circular single-stranded (ss) DNA genomes that cause devastating diseases in crops worldwide. In many geminiviruses, a small virus-encoded protein, C4, has been identified as the main symptom determinant (Medina-Puche *et al*., 2021). In the case of tomato yellow leaf curl virus (TYLCV), transgenic expression of C4 in *Arabidopsis thaliana* (hereafter referred to as Arabidopsis) or tomato is sufficient to trigger severe developmental alterations that resemble the symptoms during infection (Rosas-Diaz *et al*., 2018; Medina-Puche *et al*., 2020) (Figure S1). The ability of C4 from TYLCV to cause symptoms requires its presence at the plasma membrane, since a non-myristoylable mutant version of the protein, which accumulates in chloroplasts, does not noticeably affect plant development in Arabidopsis (Rosas-Diaz *et al*., 2018) or tomato (Figure S1).

In the geminivirus tomato yellow leaf curl Yunnan virus (TLCYnV), C4 has been proposed to cause symptoms through its nucleocytoplasmic shuttling and sequestering of the plant kinase NbSKη from the nucleus to the plasma membrane through protein-protein interaction (Mei *et al*., 2018a; Mei *et al*., 2018b). C4 from TYLCV, however, does not interact with NbSKη (Figure S2a), suggesting that this is not a conserved strategy and that a different molecular mechanism must be at play in this case. We reasoned that this effect of C4 from TYLCV would most likely be based on the physical interaction with another plant protein; with the aim to identify such a hypothetical protein, a yeast two-hybrid (Y2H) screen using C4 as bait against a TYLCV-infected tomato library was performed (Rosas-Diaz *et al*., 2018). This screen unveiled the tomato orthologues of four members of a plant-specific protein family previously described in Arabidopsis, RCC1-like domain-containing (RLD) proteins RLD1-4 (Furutani *et al*., 2020), which we have named SlRLD1 and SlRLD2, as interactors of C4 (Table S1). The RLD family in Arabidopsis comprises eight members, RLD1-8, harbouring conserved protein domains, including the namesake RCC1-like domain and a BREVIS RADIX (BRX) domain (Briggs *et al*., 2006) close to the C-terminal end (Figure 1a). Of the eight RLD members, only RLD1-4, the orthologues of the tomato proteins identified as interactors of C4, are expressed in vegetative tissues (Furutani *et al*., 2020), including the vasculature (Figure 1b; Figure S2e). Independent clones isolated from the Y2H screen pointed at the BRX domain as the minimum C4-interacting domain in the RLDs (Table S1); the specific interaction between the BRX domain of the RLD proteins (BRXD) and C4 was confirmed in yeast (Figure 1c; Figure S2). RLD proteins have been previously shown to localize in endomembrane compartments, and regulate intracellular membrane trafficking (Furutani *et al*., 2020; Wang *et al*., 2022a). The gravitropic response regulatory proteins LAZY1 and LAZY1-LIKE (LZY) (Li *et al*., 2007; Yoshihara & Iino, 2007; Yoshihara *et al*., 2013) can interact with RLD proteins and recruit them to the plasma membrane, which upon gravistimulation leads to the polarized localization of the auxin transporter PIN-FORMED 3 (PIN3) (Friml *et al*., 2002), polar auxin transport, and the subsequent developmental responses (Furutani *et al*., 2020). Strikingly, co-expression of C4-GFP also led to the recruitment of RFP-RLD proteins to the plasma membrane (Figure 1d; Figure S3a). Since LZY proteins interact with the BRX domain of the RLDs (Furutani *et al*., 2020), as observed for C4, we wondered whether this viral protein might compete with the former for RLD binding. Indeed, competitive bimolecular fluorescence complementation (BiFC) as well as fluorescence resonance energy transfer (FRET)-fluorescence lifetime imaging microscopy (FLIM) assays demonstrated that C4 outcompetes LZY3 in RLD binding (Figure 1e-g; Figure S3c, d).

**Figure 1.**
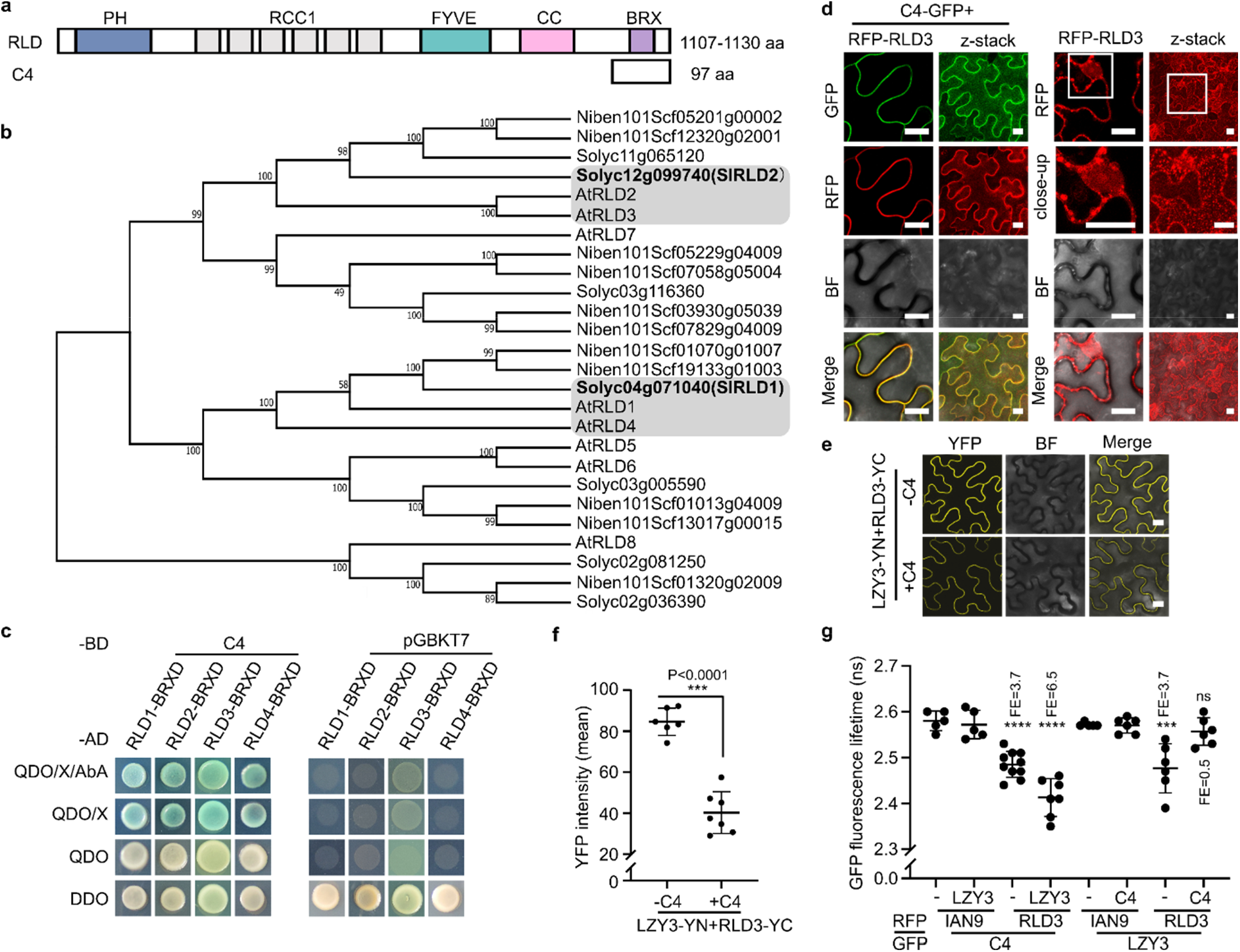
C4 interacts with RLD proteins, recruits them to the plasma membrane, and outcompetes an endogenous interacting partner. a. Domain architecture of RLDs; C4 is shown aligned with its minimal interaction domain in RLDs. PH: pleckstrin homology domain; RCC1: regulator of chromosome condensation 1-like domain; FYVE: Fab1/YGL023/Vps27/EEA1 domain; BRX: Brevis radix domain. The numbers represent the length of the corresponding proteins, in amino acids (aa). b. Phylogenetic tree of RLD proteins from Arabidopsis (AtRLDs), tomato (*Solanum lycopersicum*) (Solyc), and *N. benthamiana* (Niben101), generated by MEGA 11 and based on the full-length proteins. Bold letters indicate RLDs from tomato (SlRLDs) isolated as C4 interactors in a yeast two-hybrid TYLCV-infected tomato, and further characterized in this study. RLD proteins used in this study are contained in the grey boxes. c. Interaction between C4 and the BRX domain (BRXD) of Arabidopsis RLD1-4 by yeast two-hybrid. d. Subcellular localization of RFP-RLD3 transiently expressed in *N. benthamiana* leaves in the presence or absence of C4-GFP. White rectangles indicate close-ups. Images were taken at 2 days post-agroinfiltration (dpa). e. Interaction between LZY3 (LZY3-YN) and RLD3 (RLD3-YC) as detected by bimolecular fluorescence complementation (BiFC) upon transient expression in *N. benthamiana* leaves with or without C4. Images were taken at 2 dpa. Laser intensity was kept equal during image acquisition for all samples. f. YFP intensity of the samples in e, quantified using ImageJ. Each dot represents the YFP intensity value obtained for one technical replicate consisting of one field; lines represent the average value per sample. A minimum of six fields were analyzed per combination; asterisks indicate a statistically significant difference according to Student’s t test (***, *P*<0.001). g. Interaction between LZY3 and RLD3 in the presence or absence of C4, and interaction between C4 and RLD3 in the presence or absence of LZY3, as measured by FRET-FLIM upon transient co-expression in *N. benthamiana* leaves. Samples were taken at 2 dpa. The membrane protein IAN9 is used as negative control. FE, FRET efficiency. Each dot represents the GFP fluorescence lifetime (ns, nanoseconds) obtained for one technical replicate consisting of one field; lines represent the average value per sample. Significant differences between groups were determined by one-way ANOVA (*P*<0.0001, F=23.76, df=7), followed by multiple comparisons of means by applying Tukey test; asterisks represent statistically significant differences at: ****, *P*< 0.0001; ***, *P*< 0,005; ns, not significant. Error bars represent standard deviation. The scale bar is 20 μm in d and e. BF, bright field. Z-stack shows the maximum projection of a vertical cross-section through the observed cells. The experiments in c, d, e and g were performed three times with similar results; one representative replicate is shown here. C4, C4 from TYLCV.

A 14-aa domain named CCL present in LZY proteins mediates their interaction with the BRX domain in the RLDs (Furutani *et al*., 2020). Interestingly, inspection of the C4 protein sequence led to the identification of a similar domain, which we called CCL-like (Figure 2a). This domain is present in C4 from TYLCV and conserved in multiple viruses belonging to the same geminivirus genus, *Begomovirus* (Figure S4), but absent in the closely related tomato yellow leaf curl virus-Mild (TYLCV-Mild; Morilla *et al*., 2005) (Figures S4, S5), which produces milder symptoms. Of note, the C4 protein from TYLCV-Mild does not interact with the RLDs in yeast (Figure S5). In order to evaluate the relevance of the CCL-like domain for the interaction with the RLDs, we generated two different mutants, named CCLm1 and CCLm2, in which we replaced seven conserved residues in the CCL-like domain by alanine, or this stretch of amino acids with those present in the C4 protein from TYLCV-Mild, respectively (Figure 2a). For both mutants, the interaction with RLD proteins was abolished or largely reduced, as detected by Y2H and FRET-FLIM, and they were unable to recruit RFP-RLD proteins to the plasma membrane and to outcompete LZY3 (Figure 2b-d; Figures S6, S7). C4_CCLm1_ and C4_CCLm2_, however, localized at the plasma membrane, like the wild-type C4, and retained the ability to associate with previously described interacting partners of C4, namely the plasma membrane receptor kinases BARELY ANY MERISTEM 1 (BAM1) and CLAVATA 1 (CLV1), in Y2H, FRET-FLIM, and co-immunoprecipitation (co-IP) assays, indicating that general protein structure or stability are not affected by these mutations (Figure 2e, f; Figure S7). Importantly, and in agreement with the previously suggested function of the C4-BAM1 interaction (Rosas-Diaz *et al*., 2018; Fan *et al*., 2021), the cell-to-cell spread of silencing in the reporter SUC:SUL plants (Himber *et al*., 2003) was reduced by expression of C4_CCLm1_ and C4_CCLm2_ (Figure 2e-h; Figure S8). The CCL-like motif is required but not sufficient to mediate the interaction with RLDs, since a mutant version of C4 from TYLCV-Mild, in which the amino acids in positions 52-62 are replaced with those present in the TYLCV orthologue, hence adding a CCL-like motif, does not acquire the ability to interact with RLDs (Figure S5). These results suggest that the CCL-like domain present in geminiviral C4 proteins has evolved as a mimic of the CCL domain present in the host LZY proteins and enables the specific interaction with and recruitment of RLD proteins.

**Figure 2.**
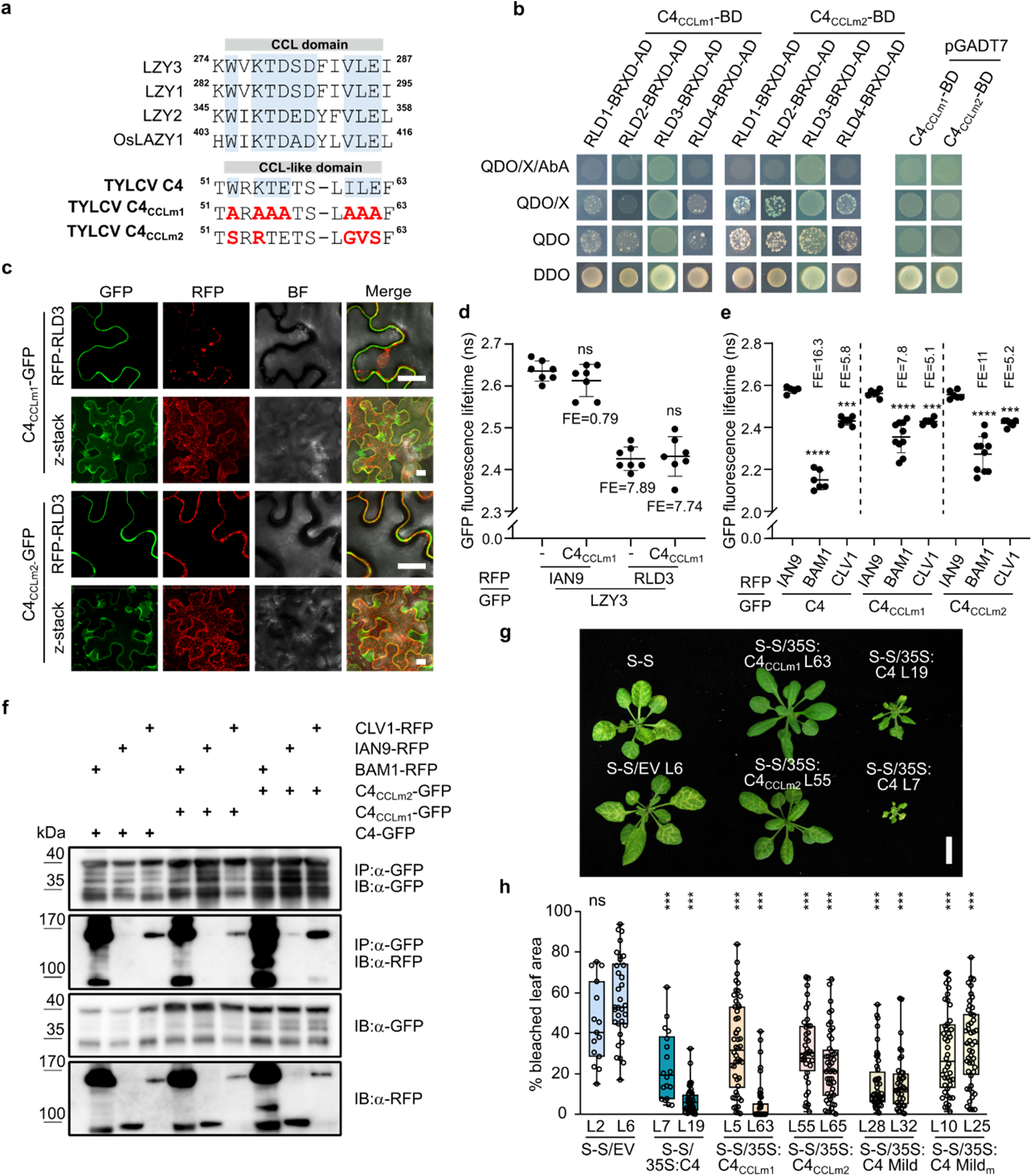
C4 specifically interacts with RLD proteins through a host-mimicking CCL-like domain. a. Alignment of amino acid sequences of LZY1, LZY2, LZY3, and OsLAZY1 in the region containing the CCL domain (Furutani *et al*., 2020). Below, identification of the CCL-like domain in C4 from TYLCV and design of C4_CCLm1_ and C4_CCLm2_ mutants; amino acid substitutions relative to the original C4 sequence are indicated in bold red letters. Numbers indicate the amino acid position in the corresponding proteins. b. Analysis of the interaction between C4_CCLm1_/C4_CCLm2_ and the BRX domain (BRXD) of Arabidopsis RLD1-4 by Y2H. c. Co-expression of RFP-RLD3 and C4_CCLm1_/C4_CCLm2_-GFP in *N. benthamiana* leaves. Images were taken at 2 days post-agroinfiltration (dpa). BF, bright field. Z-stack shows the maximum projection of the vertical cross-sections obtained through the observed cells. Scale bar, 20 μm. d. Interaction between LZY3 and RLD3 with or without C4_CCLm1_, measured by FRET-FLIM upon transient co-expression in *N. benthamiana* leaves. FE, FRET efficiency. e, f. Interaction between C4, C4_CCLm1_, or C4_CCLm2_ and the plasma membrane-localized receptor kinases BAM1 or CLV1 by FRET-FLIM (e) and co-IP (f) upon transient co-expression in *N. benthamiana* leaves. In d, e, f, samples were taken at 2 dpa; the membrane protein IAN9 is used as negative control, and the interaction between C4 and BAM1 is used as positive control. FE, FRET efficiency. In d, e, each dot represents the GFP fluorescence lifetime (ns, nanoseconds) obtained for one technical replicate consisting of one field; lines represent the average value per sample; and error bars correspond to standard deviations. Significant differences between groups were determined by one-way ANOVA (for d, *P*<0.0001, F=70.21, df= 3; for e, *P*<0.0001, F=51.63, df=8), followed by multiple comparisons of means by applying Tukey test; asterisks represent statistically significant differences at: ****, *P*<0.0001; ***, *P*<0.005; ns, not significant. IP: immunoprecipitation; IB: immunoblotting. The experiments in b, c and d were performed three times with similar results. The experiments in e and f were performed two times with similar results. One representative replicate is shown here. g. Representative phenotypes of bleaching suppression in 4-week-old SUC:SUL (S-S) Arabidopsis rosettes expressing C4, or the mutant forms C4_CCLm1_ or C4_CCLm2_ (T2 generation). Scale bar, 2 cm. S-S/EV rosettes (from plants transformed with the empty vector) are included as control. h. Bleaching quantification in different lines of 4-week-old SUC:SUL (S-S) Arabidopsis plants expressing C4 from TYLCV (C4), C4 from TYLCV-Mild (C4 Mild) or their respective mutant forms (T2 generation). In the box and whiskers graph, each dot represents bleaching values (%) of an individual leaf, and error bars indicate the highest/lowest values. Three leaves per rosette, and a minimum of ten rosettes per line, were analyzed, when possible. Differences between groups were assessed by applying Kruskal-Wallis test (*P*=4.62E-40, H=215.43, df =11), followed by pairwise comparisons between each group and the reference group (S-S/EV line 6) with Mann-Whitney U test (significance 0.0045, after Bonferroni’s correction for multiple comparisons); asterisks represent statistically significant differences at: ***, *P*<0.001; ns, not significant. C4, C4 from TYLCV; C4 Mild, C4 from TYLCV-Mild; S-S, SUC:SUL plants; EV, empty vector-transformed S-S plants.

Strikingly, transgenically expressed C4_CCLm1_ and C4_CCLm2_ in Arabidopsis did not cause any obvious phenotype, as opposed to expression of the wild-type C4 (Figure 3a, Figure S10), despite reaching similar expression levels (Figures S9, S10), indicating that an intact CCL-like domain is required for C4 to trigger developmental alterations. Supporting the idea that the interaction between the CCL-like domain in C4 and the RLDs mediates the impact of the viral protein on plant development, transgenic plants expressing C4 partially phenocopy *rld* multiple mutants (Figure 3b-d), and C4-triggered alterations are alleviated by overexpression of *RLD3* (Figure 3c Figure S11). Since C4-expressing plants show altered gravitropic set point angles (GSAs) (Figure 3d; Figure S12), as *rld* mutants do (Furutani *et al*., 2020), we decided to test root gravitropic responses in these plants and those expressing the CCL-like domain deficient versions of C4. The root response to gravity, in terms of directionality of growth as well as asymmetric activity of the auxin-responsive promoter DR5 driving GFP (DR5:GFP; Friml *et al*., 2003), was found defective in plants expressing wild-type C4, but not C4_CCLm1_ or C4_CCLm2_, suggesting that the interaction between C4 and RLDs interferes with polar auxin transport in gravitropic responses (Figure 3e, f). In line with the idea that polar auxin transport is altered by C4 through its disruption of RLD function, functional enrichment analysis of genes specifically down-regulated in plants expressing C4, but not in those expressing C4_CCLm1_ or C4_CCLm2_, unveiled the “Response to auxin” gene ontology functional category as over-represented (Figure 3g; Figure S13; Tables S2-S6). C4-expressing plants, however, accumulate wild-type-like levels of the auxin IAA, and respond normally to exogenously-applied auxin (Figure S14), suggesting that it is hormone transport, and not biosynthesis, perception, or responses, that is specifically affected.

**Figure 3.**
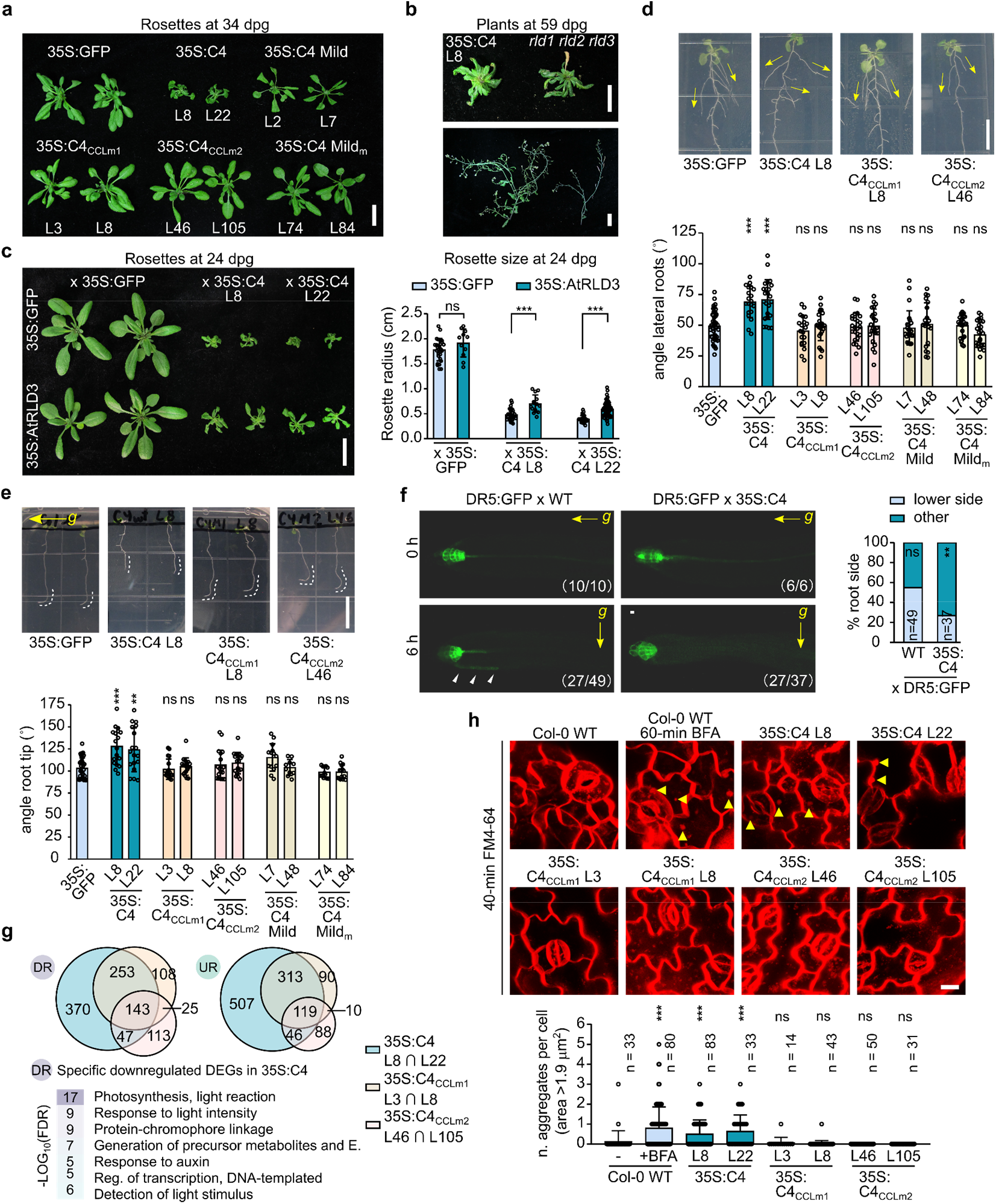
Developmental alterations triggered by stable expression of C4 from TYLCV in Arabidopsis plants rely on the interaction with the RLDs, and correlate with the interference with auxin responses and vesicle trafficking. a. Representative developmental phenotypes in Arabidopsis rosettes expressing C4 from TYLCV (C4), C4 from TYLCV-Mild (C4 Mild), or their respective mutant forms (see Figure 2a and Figure S5) (T3 generation), at 34 days post-germination (dpg). Scale bar, 2 cm. b. Comparison of developmental phenotypes between a plant expressing C4 (35S:C4 line 8, T3 generation) and a *rld1 rld2 rld3* triple mutant (Furutani *et al*., 2020). Both rosettes (upper panel) and stems with inflorescences and siliques (bottom panel) at 59 dpg are shown. Scale bar, 1 cm. c. Representative rosette phenotypes (panel on the left) and quantification of the rosette average radius (panel on the right) in the F1 generation plants resulting from crossing 35S:C4 lines 8 and 22 (T2) with 35S:AtRLD3 lines 20 and 10 (T2) parentals, respectively, at 24 dpg. In these experiments, F1 plants coming from crossing a stable line expressing GFP (35S:GFP) with 35S:AtRLD3 T2 parentals or with another 35S:GFP are used as controls. In the bar graph, each dot represents the average of three radius measurements per rosette, and error bars indicate standard deviations. A minimum of 11 rosettes per genotype were analyzed, when possible. Differences between pairs of groups were assessed by applying Mann-Whitney U test; asterisks represent statistically significant differences at: ***, *P*<0.001; ns, not significant. This experiment was repeated twice, with F1 progenies coming from two different crosses carried out independently. Scale bar, 1 cm. d. Upper pictures show representative root phenotypes in plants expressing C4 or the mutant forms C4_CCLm1_ or C4_CCLm2_ (T3 generation), at 14 dpg. Scale bar, 1 cm. Note the wider angle aperture secondary roots exhibit in plants expressing the C4 protein, indicated in the figure by the yellow arrows; a stable line expressing GFP (35S:GFP) is shown as reference. Bottom panel shows quantifications of the lateral root angles relative to the main roots in plants expressing C4, C4 Mild, or their respective mutant forms (T3 generation). In the bar graph, each dot represents the angle measurements of a lateral root, and error bars indicate standard deviations. Angles of four different roots per plant were measured, and a minimum of 6 plants were analyzed per line. Significant differences between groups were determined by one-way ANOVA (*P*=2.39E-16, F=11.474, df=10), followed by multiple comparisons of means between the C4-expressing lines against the reference group (35S:GFP), by applying Dunnett’s test; asterisks represent statistically significant differences at: ***, *P*<0.001; ns, not significant. e. Upper pictures show representative root phenotypes in plants expressing C4 or the mutant forms C4_CCLm1_ or C4_CCLm2_ (T3 generation), at 7 dpg, after growing for 12 h rotated 90 degrees with respect to their initial growing position: the new *g* force direction is depicted in the pictures by the yellow arrow on the left. Scale bar, 1 cm. Note the wider aperture of the tip angle primary roots exhibit in plants expressing the C4 protein, and indicated in the figure by the white dotted lines in root tips; a stable line expressing GFP (35S:GFP) is shown as reference. Bottom panel shows quantifications of the root tip angles in plants expressing C4, C4 Mild or their respective mutant forms (T3 generation), after receiving the same treatment. In the bar graph, each dot represents the individual measurement of a root tip angle, and error bars indicate standard deviations. Root tip angles of a minimum of 14 plants were analyzed per line. Differences between groups were assessed by applying Kruskal-Wallis test (*P*=6.24E-09, H=58.75, df=10), followed by pairwise comparisons between each group and the reference group (35S:GFP) with Mann-Whitney U test (significance 0.005, after Bonferroni’s correction for multiple comparisons); asterisks represent statistically significant differences at: ***, *P*<0.001; **, *P*=0.002; ns, not significant. f. Activation of auxin responses in root tips upon changes in gravitropic stimuli. Pictures on the left show GFP fluorescence in the root tips of the F1 plants resulting from crossing the stable reporter line DR5:GFP with either C4-expressing plants (35S:C4 lines 8 and 22, T2 generation) or WT (as control), at 5 dpg, before and after 6 h of reorientation. Arrowheads indicate asymmetric distribution of GFP signal in WT (27 out of 49) after 6 h, whereas symmetric expression was detected in C4-expressing plants (27 out of 37) after the same time. The *g* force direction is depicted in the pictures by yellow arrows. In brackets, numbers of roots showing the specific phenotype over the total checked. Scale bar, 10 μm. On the right, the stacked bar graph shows the aggregate data obtained in 4 independent experiments quantifying the number of roots (%) with asymmetrical expression of GFP fluorescence (lateral) in the root tip after 6 h over the total checked (n). Statistical differences in the % distributions were assessed by applying Fisher’s Exact test; asterisks represent statistically significant differences at: **, *P*<0.01; ns, not significant. n = number of roots. g. Transcriptomic analyses performed on 12-days-old seedlings, including 35S:C4 (lines 8 and 22, T3), 35S:C4_CCLm1_ (lines 3 and 8, T3), 35S:C4_CCLm2_ (lines 46 and 105, T3), and WT as reference. Venn diagrams show the number of overlapping genes identified as down- (DR) or upregulated (UR) in each specific genotype, when compared to the reference WT. In the bottom part, the GO enrichment analysis shows GO categories specifically over-represented in the subset of DR genes in the 35S:C4 genotype: note the presence of ‘Response to auxin’ in this list. FDR, false discovery rate. h. In the upper panels, distribution of membrane-related structures stained with FM4-64 (8 μM, 40 min) in epidermal cells of Col-0 WT, 35S:C4, 35S:C4_CCLm1_, and 35S:C4_CCLm2_ transgenic lines (T3) is compared to that of Col-0 WT treated with Brefeldin A (BFA; 70 μM, 60 min). Confocal images represent the z-stack (maximum projection) of representative phenotypes. Yellow arrowheads indicate the presence of BFA bodies in BFA-treated WT cotyledons, or vesicle aggregates reminiscent of BFA bodies in C4-expressing, non-treated plants. Non-treated, FM4-64-stained WT plants are shown as control. Scale bar, 10 μm. In the bottom panel, each dot of the bar graph represents an individual count of the number of aggregates bigger than 1.9 μm^2^ found per cell, error bars indicate standard deviations, and n indicates the total number of cells analyzed; a minimum of 14 cells per line were analyzed. Differences between groups were assessed by applying Kruskal-Wallis test (*P*=1.24E-16, H=89.99, df=7), followed by pairwise comparisons between each group and the reference group (non-treated WT) with Mann-Whitney U test (significance 0.007, after Bonferroni’s correction for multiple comparisons); asterisks represent statistically significant differences at: ***, *P*<0.001; ns, not significant. C4, C4 from TYLCV; C4 Mild, C4 from TYLCV-Mild; dpg, days post-germination; Energy, E. (for GO category); ⋂, overlapping genes; BFA, Brefeldin A.

*rld* loss-of-function mutants display defective intracellular membrane trafficking, visible as appearance of Brefeldin A (BFA) body-like structures following staining with the membrane-selective dye FM4-64 (Wang *et al*., 2022a). In keeping with the apparent C4-mediated disruption of RLD function through physical interaction, epidermal cells of transgenic Arabidopsis plants expressing wild-type C4 similarly exhibit BFA body-like aggregates, as opposed to wild-type plants or plants expressing C4_CCLm1_ or C4_CCLm2_ (Figure 3h). Further supporting a causative link between impairment of RLD function and the C4-mediated developmental alterations, wild-type Arabidopsis plants subjected to long-term BFA treatment partially phenocopy C4 transgenic plants (Figure S15). This effect is more noticeable in plants expressing C4_CCLm1_ or C4_CCLm2_, where BFA treatment might be complementing the lack of a CCL-like motif, while the rest of C4 functions would be maintained, and in which downward curling of leaves can be observed (Figure S15).

In order to evaluate the contribution of the disruption of RLD function to the viral infection, we generated an infectious TYLCV clone in which the C4 protein produced presents the replacement of two residues in the CCL-like domain described as essential for the interaction with the RLDs (Furutani *et al*., 2020), W_52_ and L_61_ in C4, by serines. Of note, the amino acid sequence of the viral protein Rep, encoded by an overlapping open reading frame, is not changed in this mutant virus (Figure 4a). As expected, this C4 mutant version, which we named C4_CCLm3_, did not interact with RLDs nor recruit them to the plasma membrane (Figure S16), but, like the other CCL-like domain-deficient mutants, maintained the capacity to interact with BAM1 and CLV1 in Y2H, FRET-FLIM, and co-IP assays (Figure S17). Importantly, local infection assays allowed us to determine that the selective recruitment of RLDs to the plasma membrane also occurs in the context of the viral infection in a C4- and CCL-like domain-dependent manner (Figure 4b). Unexpectedly, given the conservation of the CCL-like domain among begomoviruses (Figure S4), the TYLCV mutant expressing C4_CCLm3_ (TYLCV^C4_CCLm3_^) accumulated to wild-type-like levels in systemic leaves of both tomato and the model *Solanaceae* species *Nicotiana benthamiana*, although symptoms were completely abolished (tomato) or reduced (*N. benthamiana*) (Figure 4c,d; Figure S17). Therefore, viral accumulation and symptom development can be uncoupled in the TYLCV-tomato pathosystem.

**Figure 4.**
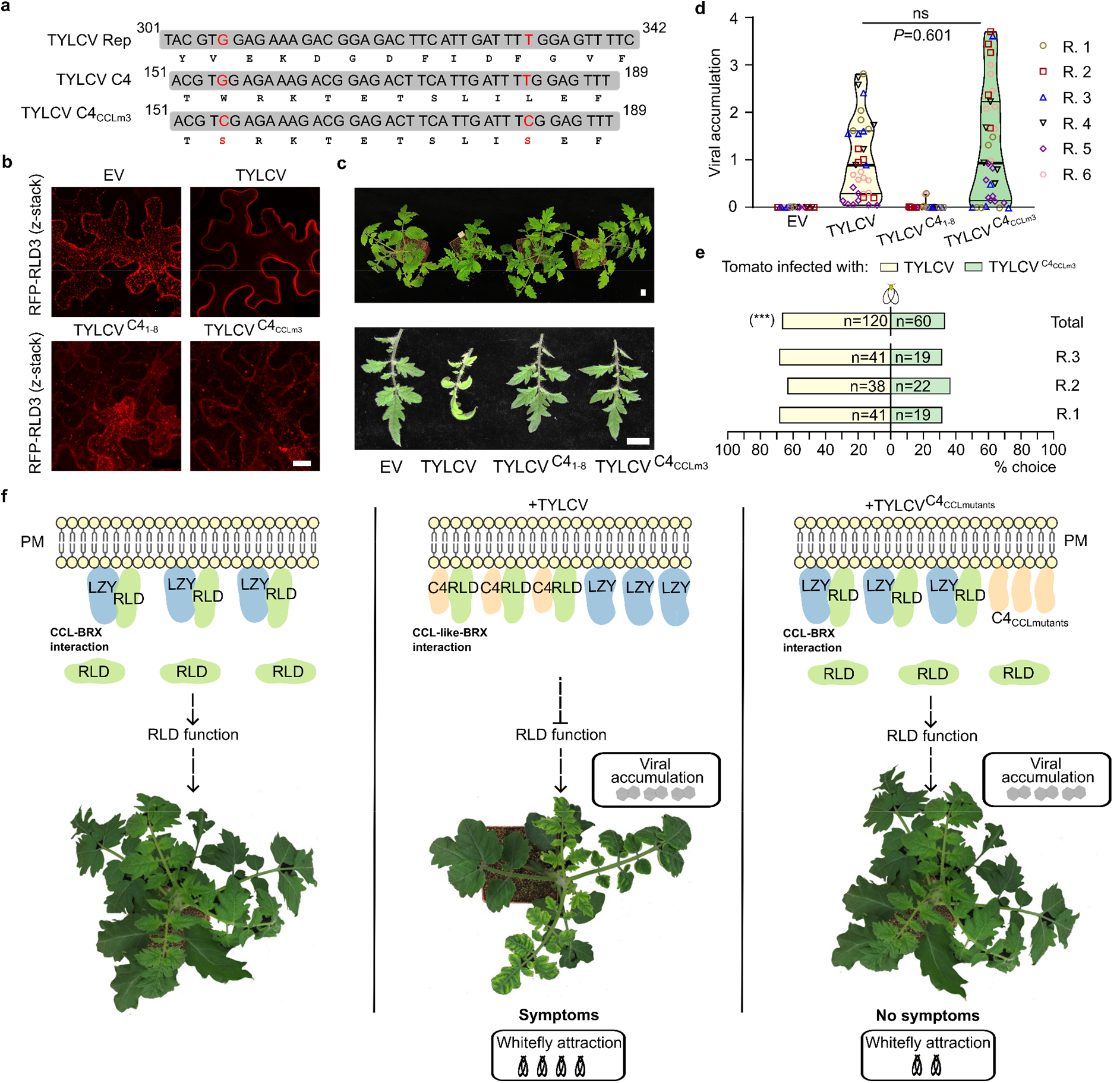
The interaction between C4 from TYLCV and RLD proteins underlies symptom development during viral infection, which is required for insect vector attraction but not for viral accumulation. a. Design of the C4_CCLm3_ mutant and resulting C4 and Rep nucleotide and amino acid sequences; changes with respect to the original sequence are indicated in red. Note that nucleotide substitutions to generate C4_CCLm3_ do not alter the Rep protein sequence. Numbers indicate nucleotide positions in the corresponding open reading frame. b. Subcellular localization of RFP-RLD3 in uninfected cells (EV, co-transformed with the empty vector), or cells inoculated with TYLCV, TYLCV^C4_1-8_^, or TYLCV^C4_CCLm3_^ infectious clones in *N. benthamiana* leaves. The images are z-stack maximum projections and were taken at 2 days post-agroinfiltration (dpa). Scale bar, 20 μm. These experiments were repeated two times with similar results; one replicate is shown here. c. Developmental phenotype of tomato plants agroinoculated with TYLCV, TYLCV^C4_1-8_^, TYLCV^C4_CCLm3_^, or empty vector (EV) as control. Images were taken at 28 days post-inoculation (dpi). Scale bar, 2 cm. d. Viral accumulation in systemic tissue of tomato plants inoculated with the viruses in c, at 28 dpi; violin plots show the aggregate results obtained in 6 independent experiments, with each dot representing one biological replicate consisting of an individual plant; thick lines represent median values and thin lines the lower/upper quartiles. A minimum of 5 plants were analyzed per virus and experiment, when possible. Differences between TYLCV and TYLCV^C4_CCLm3_^ were assessed by applying Mann-Whitney U test (*P*=0.601; ns, not significant). TYLCV^C4_1-8_^, a mutant virus carrying a premature stop codon mutation in the C4 sequence, resulting in the translation of the first 8 aa only (Rosas-Diaz *et al*., 2018), was included as control; EV, empty vector. In (d) and (e), R indicates each of the independent replicates. e. Preference shown by the TYLCV insect vector *Bemisia tabaci* (whitefly) in dual choice assays. For pairwise comparison, preference between TYLCV- or TYLCV^C4_CCLm3_^-infected tomato leaflets was recorded individually for 60 adult whiteflies at 21 dpi, and the final numbers are summarized in the horizontal bar graph. Each bar represents the % of whiteflies with the indicated preference over the total checked (% choice), with n representing the number of whiteflies. Results obtained in three independent experiments as well as the aggregate data are shown. Statistically significant differences were assessed by applying binomial test (N=180, test proportion 0.5); asterisks indicate significant differences at: \*\*\**P*<0.001. f. Model for the role of C4 in symptom development during viral infection. In the absence of infection (left panel), RLD function, which partially depends on its interaction with and recruitment by LZY proteins, contributes to normal plant development. When TYLCV infects the cell (middle panel), its C4 protein interacts with RLD proteins and recruits them to the plasma membrane (PM), to which it is associated by myristoylation. C4 outcompetes LZY proteins, and potentially other endogenous interactors of RLD proteins, for RLD binding; this results in the disruption of RLD function, and the concomitant alterations of development that are recognized as viral symptoms, which ultimately act as attractants for the whitefly insect vector. This higher whitefly attraction to virus-infected plants may favor acquisition and vector transmission of TYLCV. If plants are artificially inoculated with a mutant virus producing a CCL-like-domain-deficient version of C4 (TYLCV^C4_CCl m3_^) (right panel), the interaction with RLDs is abolished, so RLD function is maintained and plant development is not affected, rendering infected plants symptomless and reducing attractiveness to whiteflies. Of note, viral accumulation is not significantly affected by the lack of a CCL-like domain in C4, indicating that symptom development and viral performance in the plant host can be uncoupled.

Considering the high pace of evolution displayed by geminiviruses, including TYLCV (Duffy & Holmes, 2009), sequence conservation strongly argues in favour of biological relevance. Since the CCL-like motif in C4 is broadly conserved and determines symptom development, but does not seem to play a significant role in viral performance in the plant, we wondered whether it may contribute to viral transmission by the insect vector, an essential part of the viral cycle in nature. TYLCV is exclusively transmitted by the whitefly *Bemisia tabaci* (Navot *et al*., 1991), which was recently shown to be preferentially attracted to TYLCV-infected compared to uninfected tomato plants based on the perception of visual cues (Ontiveros *et al*., 2022). In agreement with these results, choice assays (Figure S18) demonstrated that *B. tabaci* displays a preference for tomato plants infected with TYLCV (symptomatic) versus those infected with TYLCV^C4_CCLm3_^ (asymptomatic), even when viral load is comparable (Figure 4e; Figure S18). It can therefore be concluded that the symptoms of TYLCV infection, which are triggered by C4 through the CCL-like domain-dependent interference with RLD function, are not determinant of viral replication and *in planta* movement, but promote insect vector attraction and hence ultimately viral spread (Figure 4f). Interestingly, the C4 protein from TLCYnV, which triggers symptom appearance through the physical interaction with NbSKη, does not possess a CCL-like domain, which raises the idea that geminiviral C4 proteins might have evolved independent strategies to manipulate plant development for insect vector attraction.

In summary, our results demonstrate that, in the TYLCV-tomato pathosystem, symptom development follows the interaction disease model, and depends on the protein-protein interaction between a host-mimicking domain in the viral C4 and the plant RLDs. This physical association leads to the recruitment of the RLDs to the plasma membrane and the disruption of their function, including the regulation of intracellular membrane trafficking and polar auxin transport, and ultimately determines the appearance of symptoms during viral infection independently of the accumulation of the virus. Importantly, the ability of C4 to interfere with plant development is independent of its other previously described virulence functions (Luna *et al*., 2012; Rosas-Diaz *et al*., 2018; Medina-Puche *et al*., 2020), which strongly argues for specific selection. Symptoms render plants more attractive to the insect vector, hence favouring viral spread. Therefore, in this pathosystem symptoms seem to have biological relevance *per se*, and the ability of the virus to trigger them is most likely under direct selective pressure. Considering that most plant viruses are insect-transmitted, similar principles might underlie symptom development and its biological relevance in other virus-crop combinations.

## METHODS

### Plant material and viral strains

All Arabidopsis plants used in this work are of the Columbia-0 (Col-0) ecotype. The AtRLD1p:GUS, AtRLD2p:GUS, AtRLD3p:GUS, and AtRLD4p:GUS transgenic lines and the *rld1-2 rld2-2 rld3-2* mutant are from Furutani *et al*., 2020. The 35S:C4 lines have been previously described (Rosas-Diaz *et al*., 2018). The *superroot 1-1* (*sur1-1*) mutant is described in Boerjan *et al*. (1995). To generate the 35S:C4_CCLm1_, 35S:C4_CCLm2_, 35S:C4 Mild, and 35S:C4 Mild_m_ lines, wild-type (WT) Arabidopsis plants were transformed with pGWB502-C4_CCLm1_, pGWB502-C4_CCLm2_, pGWB502-C4 Mild, and pGWB502-C4 Mild_m_ constructs, respectively (see Plasmids and cloning).

Tomato plants used in this work are *Solanum lycopersicum* var. Moneymaker. To generate the 35S:C4 and 35S:C4_G2A_ lines, tomato plants were transformed with pGWB2-C4 and pGWB2-C4_G2A_ constructs, respectively (Rosas-Diaz *et al*., 2018). As control, empty vector (EV) transformed tomato plants were generated in parallel.

Plant material used in this study is summarized in Table S8.

*N. benthamiana* and tomato plants were grown in a controlled growth chamber under long-day conditions (LD, 16 h of light/8 h of dark) at 25 °C. Arabidopsis plants were grown in a controlled growth room under long-day conditions (16 h light/8 h dark) at 22°C. For *in vitro* culture, Arabidopsis seeds were surface-sterilized, sown on ½ MS medium containing 1% sucrose and 1% agar (pH5.7, KOH), and stratified for three days at 4°C in the dark, after which they were grown under LD conditions.

The tomato yellow leaf curl virus-Almeria (TYLCV-Alm, Accession No. AJ489258) (Morilla *et al*., 2005) was used as template to generate the TYLCV infectious clone (Rosas-Diaz *et al*., 2018); tomato yellow leaf curl virus-Mild (TYLCV-Mild, Accession No. X76319) (Navas-Castillo *et al*., 1997) was used as template to clone ORF C4 Mild.

### Plasmids and cloning

C4-pGADT7 and C4-pGBKT7 are described in Wang *et al*., 2022b. The binary vectors to express C4, C4-GFP, IAN9-RFP, and BAM1-RFP are described in Rosas-Diaz *et al*., 2018, and that to express CLV1-RFP is described in Garnelo Gomez *et al*., 2019. The NbSKη-pGADT7 construct is from Mei *et al*., 2018b.

The C4_CCLm1_-pGADT7, C4_CCLm1_-pGBKT7, C4_CCLm2_-pGADT7, C4_CCLm2_-pGBKT7, C4 Mild_m_-pGADT7, and C4 Mild_m_-pGBKT7 were synthesized by Sangon Biotech. C4_CCLm3_-pENTRTM/D-TOPO^®^ (Thermo Scientific) was generated with the Quick-Change Lightning Site-Directed Mutagenesis Kit (Agilent Technologies).

To construct C4 Mild-pGADT7/pGBKT7, C4_CCLm3_-pGADT7/pGBKT7, AtRLD1-pGADT7/pGBKT7, AtRLD2-pGBKT7, AtRLD3-pGADT7/pGBKT7, AtRLD4-pGADT7/pGBKT7, AtRLD1-BRX domain-pGADT7/pGBKT7, AtRLD2-BRX domain-pGADT7/pGBKT7, AtRLD3-BRX domain-pGADT7/pGBKT7, AtRLD4-BRX domain-pGADT7/pGBKT7, SlRLD1-pGADT7/pGBKT7, SlRLD2-pGADT7/pGBKT7, SlRLD1-BRX domain-pGADT7/pGBKT7, SlRLD2-BRX domain-pGADT7/pGBKT7 and AtBAM1 kinase domain (678-1003aa)-pGADT7/pGBKT7, pGADT7 and pGBKT7 were digested with High-Fidelity (HF^®^) restriction endonucleases (NEB) *EcoRI* and *BamHI*, and C4 Mild, C4_CCLm3_, AtRLD1, AtRLD2, AtRLD3, AtRLD4, AtRLD1 BRX domain, AtRLD2 BRX domain, AtRL3 BRX domain, AtRLD4 BRX domain, SlRLD1, SlRLD2, SlRLD1 BRX domain, SlRLD2 BRX domain and AtBAM1 kinase domain (aa 678-1003) were amplified by PCR, and then in-fused into the pGADT7 and pGBKT7, respectively, with ClonExpress^®^ II One Step Cloning Kit.

To generate the constructs to express C4_CCLm1_-GFP, C4_CCLm2_-GFP, C4_CCLm3_-GFP, C4 Mild-GFP, C4 Mild_m_-GFP, 35S:C4_CCLm1_, 35S:C4_CCLm2_, 35S:C4 Mild, and 35S:C4 Mild_m_, the coding sequence (CDS) of C4_CCLm1_, C4_CCLm2_, C4_CCLm3_, C4 Mild, and C4 Mild_m_ were amplified by PCR and cloned into Gateway binary vectors, with a first step of cloning into pDONR^TM^/Zeo through BP reaction, and finally into the destination vectors pGWB5 and pGWB502 (Nakagawa *et al*., 2007a; Nakagawa *et al*., 2007b) through LR reaction (Thermo Scientific).

To generate the constructs to express GFP/RFP-AtRLD1, GFP/RFP-AtRLD2, GFP/RFP-AtRLD3, 35S:AtRLD3, GFP/RFP-AtRLD4, AtRLD1-YN/YC, AtRLD2-YN/YC, AtRLD3-YN/YC, AtRLD4-YN/YC, 35S:AtLZY3, AtLZY3-YN, and GFP-AtLZY3, the CDS of AtRLD1, AtRLD2, AtRLD3, AtRLD4, and AtLZY3 were amplified by PCR and first cloned into pDONR^TM^/Zeo through BP reaction, the sub-cloned to the destination vectors pGWB506, pGWB555, pGWB502, pGTQL1211YN, or pGTQL1221YC (Nakagawa *et al*., 2007a; Nakagawa *et al*., 2007b; Lu *et al*., 2010) through LR reaction (Thermo Scientific).

The TYLCV infectious clone and its mutant version carrying a premature stop codon in the C4 gene (TYLCV^C4_1-8_^) are described in Rosas-Diaz *et al*., 2018. To generate the TYLCV^C4_CCLm3_^ mutant virus, mutations G155C (W52S) and T182C (L61S) (Figure 4a), which do not alter the protein sequence of Rep, were introduced in the coding sequence of C4 with the Quick-Change Lightning Site-Directed Mutagenesis Kit (Agilent Technologies).

All primers used in this study are listed in Table S9.

### Transient expression and viral infection assays in *N. benthamiana* and tomato

Transient expression assays were carried out as described previously with minor modifications (Wang *et al*., 2017). In brief, the *A. tumefaciens* strain GV3101 carrying the corresponding construct was liquid-cultured in LB with appropriate antibiotics at 28°C overnight. Bacterial cultures were then centrifuged at 4,000x g for 10 min and resuspended in infiltration buffer (10 mM MgCl_2_, 10 mM MES pH 5.6, 150 μM acetosyringone) to an OD_600_ = 0.5-1. Finally, Agrobacterium suspensions were placed at room temperature in darkness for at least 2 hours, and then infiltrated into the abaxial side of fully expanded young leaves of three-to-four-week-old *N. benthamiana* plants with a 1 mL needleless syringe. For experiments that needed co-expression, Agrobacterium suspensions carrying different constructs were mixed before infiltration at a 1:1 ratio. In the competitive BiFC assays, the OD_600_ of Agrobacterium suspensions carrying LZY3-YN, RLD3-YC, and C4 constructs was 0.3, 0.2, and 0.2, respectively.

Systemic viral infections were carried out as in Wu *et al*., 2021. In short, for *N. benthamiana* and tomato, the Agrobacterium cells carrying the empty vector (EV) as a negative control, or the TYLCV, TYLCV^C4_1-8_^, or TYLCV^C4_CCLm3_^ infectious clones were initially prepared as described before, and resuspended in infiltration buffer to a final OD_600_ = 0.5. Bacterial solutions were then injected in the stem of two-week-old *N. benthamiana* and three to four-week-old tomato plants (for systemic infection assays) or infiltrated into the abaxial side of fully expanded young leaves of four-week-old *N. benthamiana* plants (for local infection assays). To analyze local viral accumulation, as a proxy for replication, samples from agroinfiltrated leaves of *N. benthamiana* plants were collected at 2 days post-inoculation (dpi). To analyze systemic viral accumulation, the three youngest apical leaves from inoculated *N. benthamiana* or tomato plants were collected at 21-28 dpi.

### Generation of transgenic plants

Transformation of Arabidopsis plants, in either the WT or the SUC:SUL (S-S) (Himber *et al*., 2003) backgrounds, was performed through the floral dip method (Clough & Bent, 1998). In brief, the *A. tumefaciens* strain GV3101 carrying the corresponding construct was liquid-cultured in LB liquid medium with appropriate antibiotics at 28°C overnight. Then, the bacterial culture was pelleted by centrifugation at room temperature at 4000x g for 10 min, and resuspended in transformation solution (5% sucrose and 0.02% Silwet L-77). Inflorescences of 3-4-week-old plants were gently immersed in the bacterial suspension for 10-20 seconds, and the treated plants were wrapped in plastic film to maintain high humidity, and maintained in darkness for 16-24 hours. Finally, these plants were returned to normal (LD) growth conditions, their seeds recovered, and rounds of selection and propagation were performed until reaching the specified generation.

### Isolation of nucleic acids and quantitative PCR

In all cases, samples were taken per duplicate from apical, young leaves (for *N. benthamiana* and tomato) or rosette leaves (for Arabidopsis), and nucleic acids were extracted by using Plant RNA kit (Omega) for total RNA, or CTAB 2X solution for DNA (Murray & Thompson, 1980). Total RNA was reverse-transcribed by using iScript TM cDNA synthesis Kit (Bio-Rad) in a volume of 20 μL, and the resulting cDNA was diluted to a final volume of 200 μL. For DNA, once purified, samples were diluted 1/1000 before analysis. RT-qPCR (cDNA) or qPCR (DNA) were performed in a C1000 Touch Thermal Cycler (Bio-Rad), with PCR mixtures containing 6 μL of diluted cDNA/DNA, 1 μL of each primer (10 μM), 2 μL of water, and 10 μL of Hieff qPCR SYBR Green Master Mix (Yeasen), with the following program: 3 min at 95 °C, and 40 cycles consisting of 15 s at 95 °C, 30 s at 60 °C. As normalizer, *ACTIN (ACT2*) was used for RT-qPCR in Arabidopsis, while 25S ribosomal DNA interspacer (ITS) was used for qPCR in *N. benthamiana* or tomato samples (Mason *et al*., 2008) (see Table S9). Comparative analyses of transcripts and viral accumulation were performed by applying the 2^-ΔΔCt^ method.

### Confocal imaging

Confocal imaging was performed on a Leica TCS SP8 confocal microscope or an SMD FLCS point scanning confocal microscope (Leica Microsystems) using the pre-set sequential scanning settings for GFP with excitation (Ex):488 nm, emission (Em):500–550 nm, for RFP with Ex:561 nm, Em:570-620 nm, and for YFP with Ex: 514 nm, Em: 525-575 nm. For aniline blue staining, settings were Ex: 405 nm, Em: 448–525 nm with sequential scanning when combined with other fluorophores; for FM4-64 staining, Ex:580 nm, Em:600 to 660 nm. Z-maximum projections were generated with LAS X software.

For competitive BiFC assays, laser intensity was fixed during image acquisition throughout the experiment. Images were transformed to 8-bit by FIJI, and background noise was removed by applying default threshold settings for each image; final quantification values correspond to YFP intensity.

In Förster resonance energy transfer by fluorescence lifetime imaging (FRET-FLIM) assays, donor proteins were cloned into pGWB5 or pGWB506 (Nakagawa *et al*., 2007a; Nakagawa *et al*., 2007b) (fused to GFP), and acceptor proteins were cloned into pGWB555 (Nakagawa *et al*., 2007b) (fused to RFP). For competitive FRET-FLIM assays, potential competing proteins were cloned into pGWB2 or pGWB502 (Nakagawa *et al*., 2007a; Nakagawa *et al*., 2007b) (no tag). FRET-FLIM experiments were performed on a Leica TCS SMD FLCS confocal microscope using excitation with WLL (white light laser) and emission collected by a SMD SPAD (single photon-sensitive avalanche photodiodes) detector, as described in Rosas-Diaz *et al*., 2018. Leaf discs of *N. benthamiana* plants transiently co-expressing the proteins of interest were visualized two days after agroinfiltration.

### Yeast two-hybrid (Y2H) assay

All yeast constructs were transformed into the *Saccharomyces cerevisiae* Y2H Gold strain (Clontech) using Yeastmaker^™^ Yeast Transformation System 2 (Clontech). Transformants were grown on minimal synthetic defined (SD) media without leucine and tryptophan plates (double dropout medium, DDO) and re-suspended in 20 μL DDO liquid medium. 3-5 μL of each suspension were placed onto SD media without leucine, tryptophan, histidine, and adenine (quadruple dropout medium QDO), QDO with X-α-gal (QDO/X) and QDO/X with Aureobasidin A (QDO/X/AbA). Plates were incubated in the dark at 28°C and photographed five to seven days later. The prey plasmid pGADT7-T, encoding the SV40 large T-antigen fused with the GAL4 activation domain (AD), and the bait plasmid pGBKT7-p53, encoding murine p53 fused with the GAL4 DNA-binding domain (BD), were used as positive control. pGADT7 (AD) and pGBKT7 (BD) empty vectors were used as negative control. All yeast media were purchased from Clontech.

### Protein extraction and co-immunoprecipitation (co-IP) assays

Two days after infiltration, 0.75–1 g of infiltrated *N. benthamiana* leaves were harvested. Protein extraction, co-immunoprecipitation (co-IP), and western blot were performed as described in Rosas-Diaz *et al*., 2018. The following primary and secondary antibodies were used for western blot: mouse anti-green fluorescent protein (GFP) (M0802-3a, Abiocode, Agoura Hills, CA, USA) (1:10,000); rat anti-red fluorescent protein (RFP) (5F8, Chromotek, Planegg-Martinsried, Germany) (1:10,000); goat polyclonal anti-mouse coupled to horseradish peroxidase (Sigma, St. Louis, MO, USA) (1:15,000); and goat polyclonal anti-rat coupled to horseradish peroxidase (Abcam, Cambridge, UK) (1:15,000).

### Tissue staining and chemical treatments

For β-glucuronidase (GUS) staining, 3-day-old transgenic seedlings carrying the reporter constructs (RLD promoter:GUS) were immersed in GUS staining solution and vacuum-infiltrated for five minutes, twice. Then, samples were incubated overnight at 37°C. The following day, the staining solution was removed and the samples were washed with 75% ethanol until tissue cleared. Images were taken using Zeiss Imager M2 microscope.

For aniline blue staining, a 0.05% solution (w/v in water) of the dye was infiltrated into the abaxial side of *N. benthamiana* leaves and incubated for 30 min before imaging.

Counts of BFA-like bodies were performed in 5-day-old seedlings as described previously (Wang *et al*., 2022a). In brief, WT and transgenic Arabidopsis plants expressing TYLCV C4 (C4) or its mutant forms C4_CCLm1_ and C4_CCLm2_ (T3 generation) were pre-incubated for 40 min with 8 μM FM4-64, and then treated for 60 min with 70 μM Brefeldin A (BFA) or DMSO (control plants). Z-stack maximum projection images of stained, treated cotyledons were obtained, and counts of vesicle aggregates and their area were determined with FIJI. The average size of the aggregates per cell was determined as of 1.9 μm^2^, and the number of aggregates with areas bigger than this average was registered.

To assess the effect of prolonged BFA treatment on plant development, WT and transgenic Arabidopsis plants expressing TYLCV C4 (C4) or its mutant forms C4_CCLm1_ and C4_CCLm2_ were initially grown in vertical ½ MS plates and, at 5 days post-germination, transferred to new ½ MS plates supplemented with BFA (1, 10, or 25 mM dissolved in DMSO) or DMSO (control). Pictures of representative plants were taken after a 7-day treatment.

### Visual analysis of RNA silencing spread in Arabidopsis SUC:SUL plants

TYLCV C4 (C4), TYLCV-Mild C4 (C4 Mild), and their respective mutant forms, were expressed in the SUC:SUL background (Himber *et al*., 2003) in order to assess their effect on the cell-to-cell movement of RNA silencing. SUC:SUL plants were transformed with the appropriate constructs (see Table S8) by using the floral dip method, and 20 lines per construct were selected on hygromycin plates. At 4-5 weeks after germination, the lines showing evident reduction of bleaching were taken to the T2 generation. These plants were selected on hygromycin plates before being transferred to soil at 12 days after germination. Three weeks later, each line was sampled and the expression levels of both the endogenous *SUC2* gene and the transgenic SUC:SUL cassette were assessed by RT-qPCR (see Table S9). After confirmation of maintained expression of *SUC2*/SUC:SUL, three medium-size leaves per rosette, and a minimum of 10 rosettes per line, were photographed; bleached area was quantified with FIJI and expressed as % of total leaf area (Rosas-Diaz *et al*., 2018).

### IAA treatment and quantification of IAA content

TYLCV C4-expressing Arabidopsis plants (lines 5 and 7, T3) were grown in vertical ½ MS plates supplemented with IAA (10, 20, 30, or 40 nM IAA dissolved in 0.1% ethanol) or 0.1% ethanol (as control). At 12 days post-germination (dpg), root length was measured with FIJI, and the number of lateral roots was counted and expressed as the number of initials per cm of primary root. At least 10 plants were analyzed per replicate, and the experiment was repeated twice with similar results.

IAA content in seedlings expressing either TYLCV C4 or its mutant forms C4_CCLm1_ or C4_CCLm2_ was determined at 11 dpg. Three biological replicates (50 mg fresh weight per replicate) were retched (5mm ceramic ball; 30 sec) and successively extracted with 200 μL 80% MeOH (containing the isotopic standard 200 nM D7-IAA), 200 μL 80% MeOH, and 400 μL H2O 0,1% formic acid. 400 μL of the united fractions were partitioned against 200 μL chloroform (10 min US RT) and the upper phase was directly measured using targeted LCMS analysis. The whole extraction process was below 10°C including 5 min sonication and centrifugation steps (18,600x g).

The LCMS profiling analysis was performed using a Micro-LC M5 (Trap and Elute) and a QTRAP6500+ (Sciex) operated in MRM mode (MRMs IAA (1) quantifier ion (m/z) Q1/Q3 176.1/130, declustering potential DP 40 V, collision energy CE 20V; IAA (2) (m/z) Q1/Q3 176.1/130, DP 100 V, CE 30V; D7-IAA (1) quantifier ion (m/z) Q1/Q3 183.1/109, DP 40, CE 50 V; D7-IAA (2) (m/z) Q1/Q3 183.1/136), DP 40, CE 20 V. The dwell time for all MRMs was 10 msec. Chromatographic separation was achieved on a Luna Omega Polar C18 column (3 μm; 100 Å; 150×0.5 mm; Phenomenex) and a Luna C18(2) trap column (5 μm; 100 Å; 20×0.5 mm; Phenomenex) with a column temperature of 55 °C. The following binary gradient was applied for the main column at a flow rate of 28 μL min-1: 0 - 0.2 min, isocratic 90% A; 0.2 - 2 min, linear from 90% A to 30% A; 2 - 4.5 min, linear from 30% A to 10% A; 4.5 - 5 min, linear from 10% A to 5% A; 5 - 5.3 min, isocratic 5 % A; 5.3 - 5.5 min, linear from 5% A to 90% A; 5.5 - 6 min, isocratic 90% A (A: water, 0.1% aq. formic acid; B: acetonitrile, 0.1% aq. formic acid). The samples were concentrated on the trap column using the following conditions: flow rate 50 μL min-1: 0 - 1.5 min isocratic 95% A; 1.5 min start main gradient; 1.5 - 1.7 min isocratic 95% A. The injection volume was 50 μL. Analytes were ionized using an Optiflow Turbo V ion source equipped with a SteadySpray T micro electrode in positive (ion spray voltage: 4800 V) ion mode. The following additional instrument settings were applied: nebuliser and heater gas, nitrogen, 25 and 45 psi; curtain gas, nitrogen, 30 psi; collision gas, nitrogen, medium; source temperature, 200 °C; entrance potential, +/-10 V; collision cell exit potential, +/-25V.

The IAA content in each sample was normalized against the D7 IAA (Cambridge Isotope Laboratories) values. The over-accumulating *sur1-1* mutant was used as a positive control for the naturally occurring low IAA level in non-treated WT seedlings.

### Analysis of plant developmental phenotypes and gravitropic responses

The effect of TYLCV C4 (C4), TYLCV-Mild C4 (C4 Mild), or their respective mutant versions in root gravitropism in transgenic Arabidopsis plants (T3 generation) was evaluated as previously described (Furutani *et al*., 2020). Briefly, plants were grown in square, vertical ½ MS plates which were rotated 90 degrees at 7 days post-germination (dpg). Seedlings were then left to grow for 12 more hours, and then pictures of at least 16 seedlings were taken to quantify root tip angles with FIJI, relative to the original vertical axis, and excluding individuals where the root tip angle was lower than 90 degrees.

T2 35S:C4 Arabidopsis plants (line 9) were crossed to the DR5:GFP line (Friml *et al*., 2003); WT plants were crossed to DR5:GFP in parallel, as control. Live cell tracking was possible by using sterile chambered slides with transparent bottom parts (Lab-Tek II coverglass chamber system, no. 155361, ThermoFisher Scientific), as described in Dubreuil *et al*., 2018 and Nakamura *et al*., 2018. The F1 seeds were grown in these chambers with MS medium, placed at the interface between the MS-agar and the coverglass, allowing their direct microscope monitoring upon vertical growth. At 5 dpg, they were reoriented 90 degrees and kept in the new position for 6 hours. Finally, the chambers were placed horizontally during the imaging; GFP confocal images were obtained with a Leica TCS SP8 microscope with a 20x objective.

In order to analyze root phenotypes in C4-expressing plants at a later time, transgenic plants (T3 generation) were grown in square, vertical plates for 14 days, until lateral roots were clearly developed, and pictures of at least 16 seedlings were taken. The length of the primary roots and the angles of four lateral roots per seedling, at four different heights relative to the primary root, were measured with FIJI; the total number of lateral roots was expressed as the number of lateral roots per unit of primary root length (/cm).

35S:AtRLD3 and 35S:C4 Arabidopsis plants were generated in parallel by the floral dip method, propagated until the T2 generation, and crossed; a stable 35S:GFP plant line was crossed as control. The F1 generation was genotyped, double transformants were selected, and their phenotypes recorded at different developmental stages. Rosette size was quantified with FIJI, measuring rosette radios at three different angles per plant, which were expressed as the average rosette radius (cm).

### Phylogenetic analyses

Identification of the BRX domain in AtRLD1-4 and SlRLD1-2 was performed by using the NCBI conserved domain research tool (https://www.ncbi.nlm.nih.gov/orffinder/), the ExPASy Prosite database (https://prosite.expasy.org/), and the InterPro database and diagnostic tool (https://www.ebi.ac.uk/interpro/).

RLD protein sequences from Arabidopsis, *N. benthamiana*, and tomato, were subjected to multiple alignments with Clustal W, and phylogenetic trees were obtained through MEGA 11 software (maximum likelihood, 500 boostrap repetitions).

To identify C4 homologs in different begomoviruses, the reference genome of each virus was downloaded and analyzed in the NCBI ORFfinder tool (https://www.ncbi.nlm.nih.gov/orffinder/); multiple sequence alignments were conducted by using the BioEdit software.

### RNA sequencing and enrichment analyses

Arabidopsis seedlings were grown in vertical ½ MS plates and, at 12 days post-germination, aerial parts of at least 30 seedlings were excised, pooled, and frozen in liquid N_2_. Three replicates, containing biological material from three different plates, were generated for the WT (reference) and transgenic lines (T3 generation) expressing TYLCV C4 (C4), TYLCV-Mild C4 (C4 Mild), or their respective mutant forms. Total RNA was extracted with TRIzol reagent (Invitrogen), followed by two ethanolic precipitations with sodium acetate 3M pH 5.2 for 24 hours, and two additional cleaning steps with 70% cold ethanol. Purified samples were then prepared for RNA sequencing as previously described (Wu *et al*., 2019). Once the RNA samples were sequenced, paired-end reads were cleaned by Trimmomatic (v. 0.36) (Bolger *et al*., 2014). After trimming the adapter sequences, removing low quality bases, and filtering short reads, the resulting clean read pairs were retained for further analyses. The Arabidopsis reference genome was downloaded from *TAIR10*, and clean reads were mapped by using HISAT (Kim *et al*., 2015), with default parameters. The number of reads mapped to each gene was calculated with htseq-count script in HTSeq (Anders *et al*., 2015). Differential gene expression analyses were performed in EdgeR (Robinson *et al*., 2010). Genes with at least two-fold change in expression levels, and with an FDR<0.05 were considered as differentially expressed (DEGs). Lists of DEGs for up and downregulated genes were subjected to Gene Ontology (GO) enrichment by using ShinyGO (Ge *et al*., 2020), with the total list of genes for which reads were identified as background, and lists with the first 40 GO categories with FDR<0.05 were generated. Enrichment analyses in The Kyoto Encyclopaedia of Genes and Genomes (KEGG) pathways were carried out through the ShinyGO portal, and visualized through Pathview (Luo & Brouwer, 2013; Kanehisa *et al*., 2021); lists of KEGG pathways with FDR<0.05 were generated. Finally, GO lists were reduced by removing redundant low-level categories by using REVIGO ((Supek *et al*., 2011); GO database last updated on 22^nd^ March 2022), generating simplified lists of the 20 first GO categories with the lowest FDR values. For raw GO lists, see Table S5.

### Whitefly dual-choice assays

3-week-old tomato plants were agroinoculated with TYLCV WT or TYLCV^C4_CCLm3_^ infectious clones. After 3 weeks, viral accumulation was checked in systemic leaves by qPCR, and plants with similar viral accumulation levels were selected. Whitefly choice assays were performed as described by Ontiveros *et al*., 2022. Briefly, four different leaflets coming from TYLCV WT- or mutant-infected tomato plants (2x each) were distributed in a plastic cage (25×25 cm), disposed around a central release platform where one whitefly was placed, and only the first choice within a time frame of 15 min was recorded. For each experiment, a total of 60 adults of *Bemisia tabaci* Mediterranean biotype were used (with each whitefly constituting an individual biological replicate). The experimental design is shown in Figure S18.

### Statistical analysis

Statistical comparisons between pairs of means were performed either by applying Student’s t test (considering equal variances) or Welch’s t test (assuming unequal variances). When statistical multiple comparisons of means were required, one-way ANOVA followed by Dunnett’s test (comparisons of multiple groups to one reference group) or Tukey multiple range test (comparisons between multiple groups) were performed. Mann-Whitney U test was used to compare pairs of means when data distributions were not normal, and data transformation was not possible. Multiple comparisons between means of non-normally distributed data were performed by applying Kruskal-Wallis test, followed by pairwise comparisons of means with Mann-Whitney U test with the Bonferroni’s correction of significance level. Statistical comparisons of % distributions were carried out by applying Fisher’s exact test (in DR5:GFP-crossed plants and gravitropism experiments) or binomial test (in whitefly choice assays, with 0.5 as test proportion). Significance level was kept at 0.05 throughout the whole study. Statistical analyses were performed with either IBM SPSS Statistics v.20 or GraphPad Prism v.8 software.

## Supporting information

Table S1. SlRLD clones isolated from the Y2H screen.

Table S2. Summary of total RNA-seq data.

Table S3. Summary of C4 counts and RPKM in total RNA-seq data.

Table S4. Summary of DEGs relative to WT plants.

Table S5. Summary of GO enrichment for DEGs in C4-expressing plants relative to WT.

Table S6. Summary of KEGG pathways enrichment for DEGs in C4-expressing plants relative to WT.

Table S7. Viruses used in Figure S4.

Table S8. Plant material used in this work.

Table S9. Primers used in this work.

## ACKNOWLEDGEMENTS

The authors thank Alberto P Macho, Thorsten Nürnberger, Sebastian Wolf, and the GeminiTeam lab for critical reading of the manuscript, and Xinyu Jian, Bettina Stadelhofer, and Aurore Luque for excellent technical assistance. Work in RLD’s lab is partially funded by the Excellence Strategy of the German Federal and State Governments, the ERC-COG GemOmics (101044142), the DeutscheForschungsgemeinschaft (DFG, German Research foundation) (project numbers LO 2314/1-1 and SBF 1101/3, C08), and a Royal Society Newton Advance grant (NA140481 – NAF\R2\180857). Metabolite analytics were funded by the DFG (Projektnummer 442641014). EA is the recipient of a Marie Skłodowska-Curie Grant from the European Union’s Horizon 2020 Research and Innovation Program (Grant 896910-GeminiDECODER). BGG was the recipient of a President’s International Fellowship Initiative (PIFI) from the Chinese Academy of Science (CAS) (grant 2020PB0082).

## Notes

### Competing Interest Statement

The authors have declared no competing interest.

