## Supplementary material for "A plant virus causes symptoms through the deployment of a host-mimicking protein domain to attract the insect vector": Figures S1-S18

### SUPPLEMENTAL FIGURES

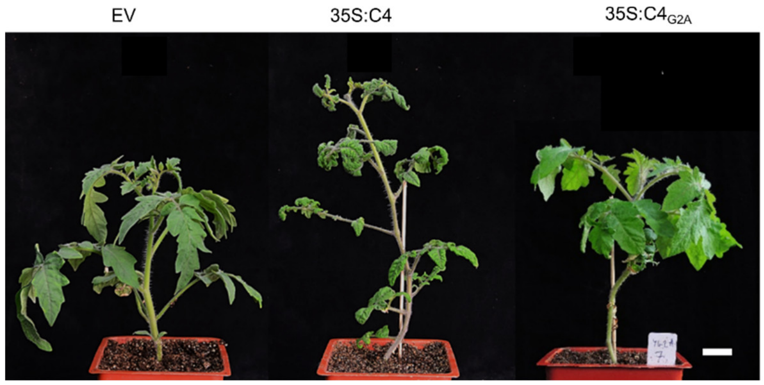


**Figure S1. C4 from TYLCV must be at the plasma membrane to trigger developmental alterations in tomato plants.**

Developmental phenotype of transgenic tomato plants expressing C4 and $C4_{G2A}$ under a 35S promoter in the T1 generation. Images were taken at 60 days post-germination. Scale bar, 2 cm.

C4, C4 from TYLCV; EV, empty vector.

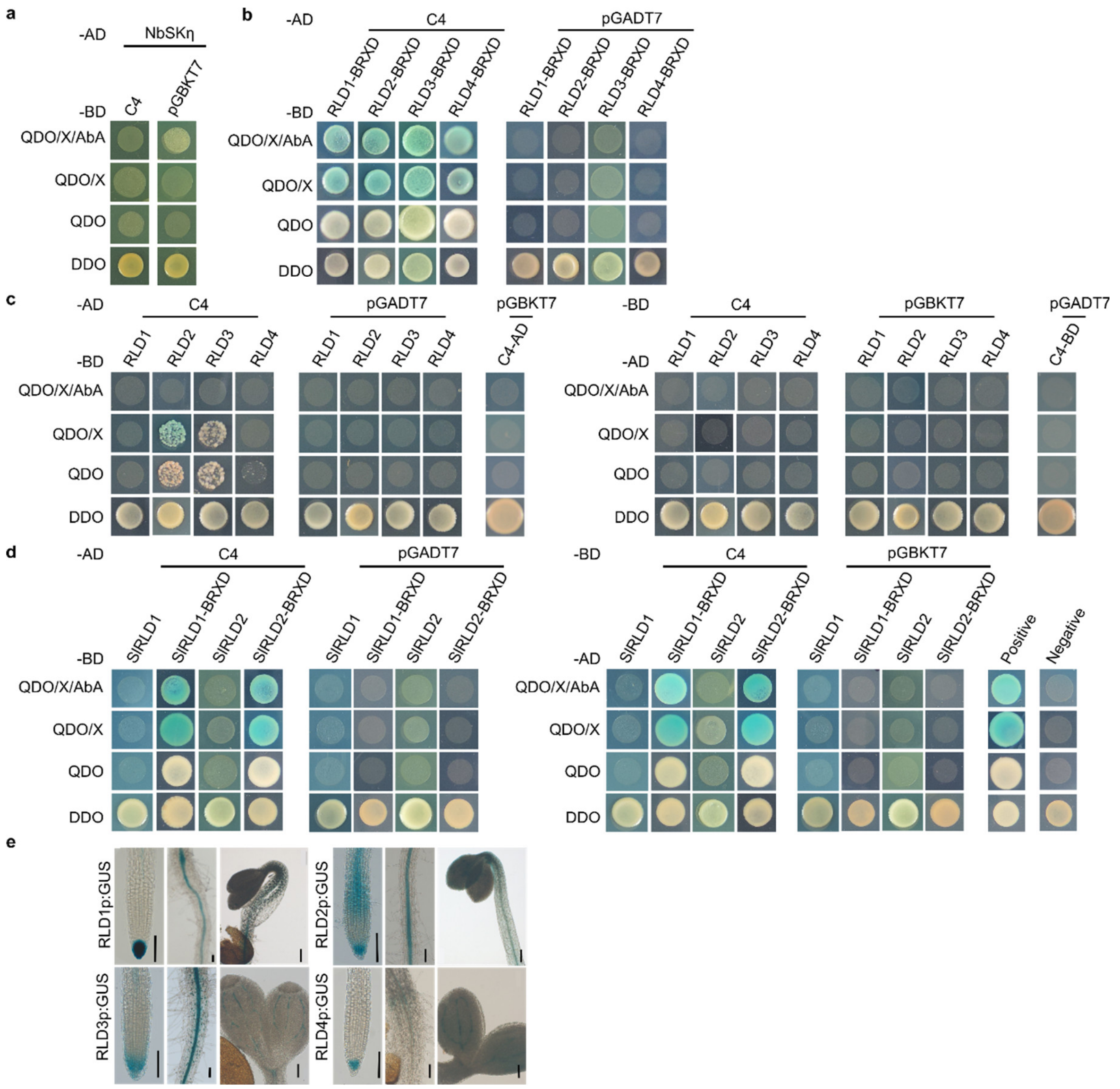


**Figure S2. C4 from TYLCV interacts with RLD proteins, of which the coding genes are expressed in the vasculature.**

a. Interaction between C4 and NbSKη, tested by yeast two-hybrid (Y2H). The NbSKη construct is from Mei *et al.*, 2018b.

b. Interaction between C4 and the BRX domain of Arabidopsis RLD1-4 (BRXD) tested by Y2H.

c. Interaction between C4 and full-length Arabidopsis RLD1-4 tested by Y2H.

d. Interaction between C4 and full-length tomato RLD1 and 2 (SlRLD1/2) or their BRX domains (BRXD).

e. Histochemical GUS staining of RLD1/RLD2/RLD3/RLD4 promoter:GUS transgenic Arabidopsis lines (T5 generation). Images were taken 3 days after germination using a Zeiss Imager M2 microscope. Three independent single-copy transgenic lines were tested per construct with similar results. Scale bar, 100 μm.

The experiments in a, b, c and d were performed three times with similar results. One replicate is shown here.

C4, C4 from TYLCV.

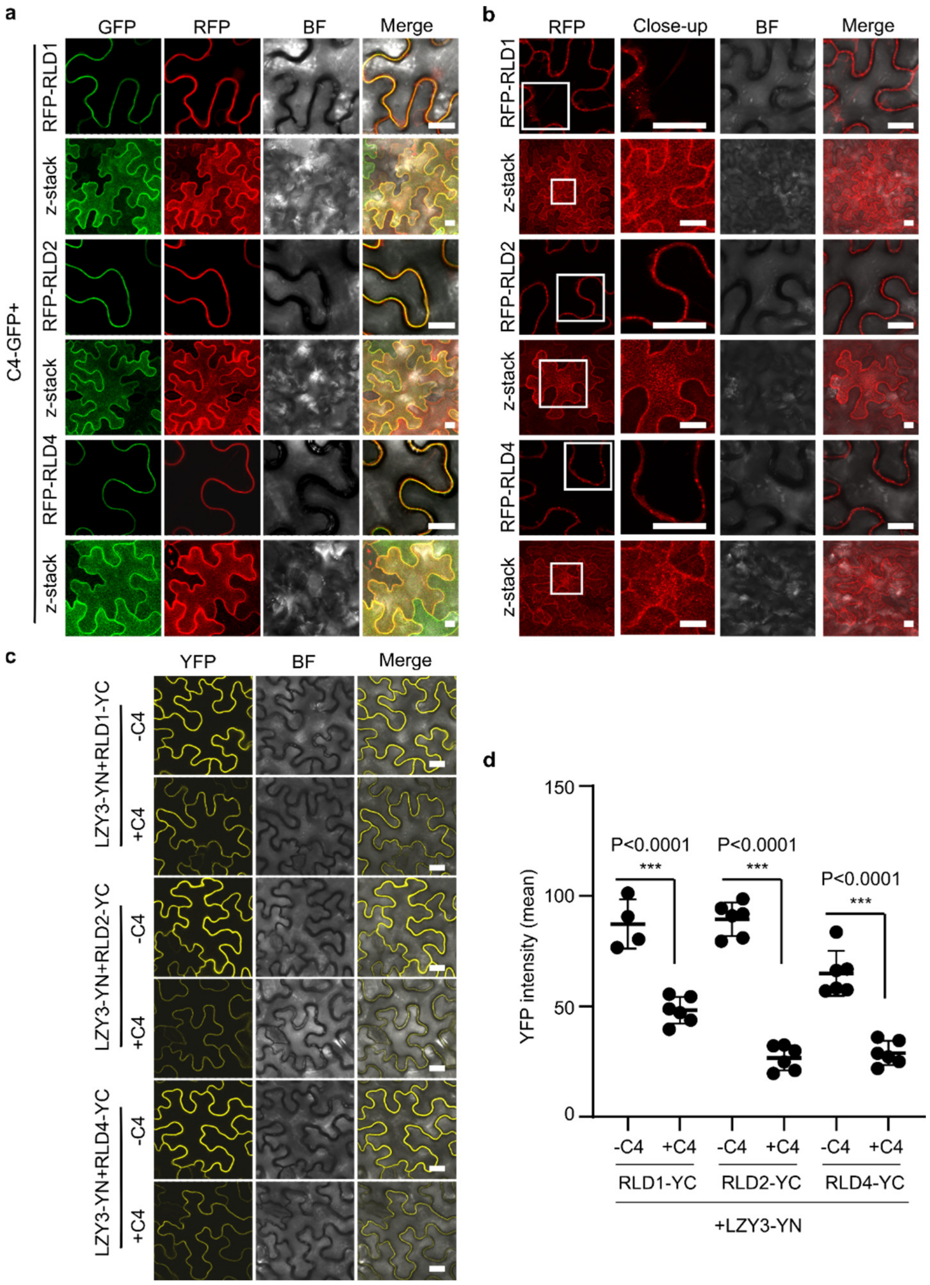


**Figure S3. C4 outcompetes LZY and recruits RLD proteins to the plasma membrane.**

a, b. Subcellular localization of RFP-RLD1, 2, and 4 transiently expressed in *N. benthamiana* leaves in the presence (a) or absence (b) of C4-GFP. White rectangles indicate close-ups. Images were taken at 2 days post-agroinfiltration (dpa).

c. Interaction between LZY3 (LZY3-YN) and RLD1, 2, an 4 (RLD1/2/4-YC) as detected by bimolecular fluorescence complementation (BiFC) upon transient expression in *N. benthamiana* leaves with or without C4. Images were taken at 2 dpa. Laser intensity was kept equal during image acquisition for all samples.

d. YFP intensity of the samples in (c), quantified using ImageJ. Each dot represents the YFP intensity value obtained for one technical replicate consisting of one field; lines represent the average value per sample. A minimum of six fields were analyzed per combination; asterisks indicate a statistically significant difference according to Student's t test (***, $P<0.001$). Images were taken at 2 dpa. BF, bright field.

Z-stack shows the maximum projection of a vertical cross-section through the observed cells. Scale bar, 20 μm. The experiments in a, b and c were performed three times with similar results. One replicate is shown here.

C4, C4 from TYLCV.

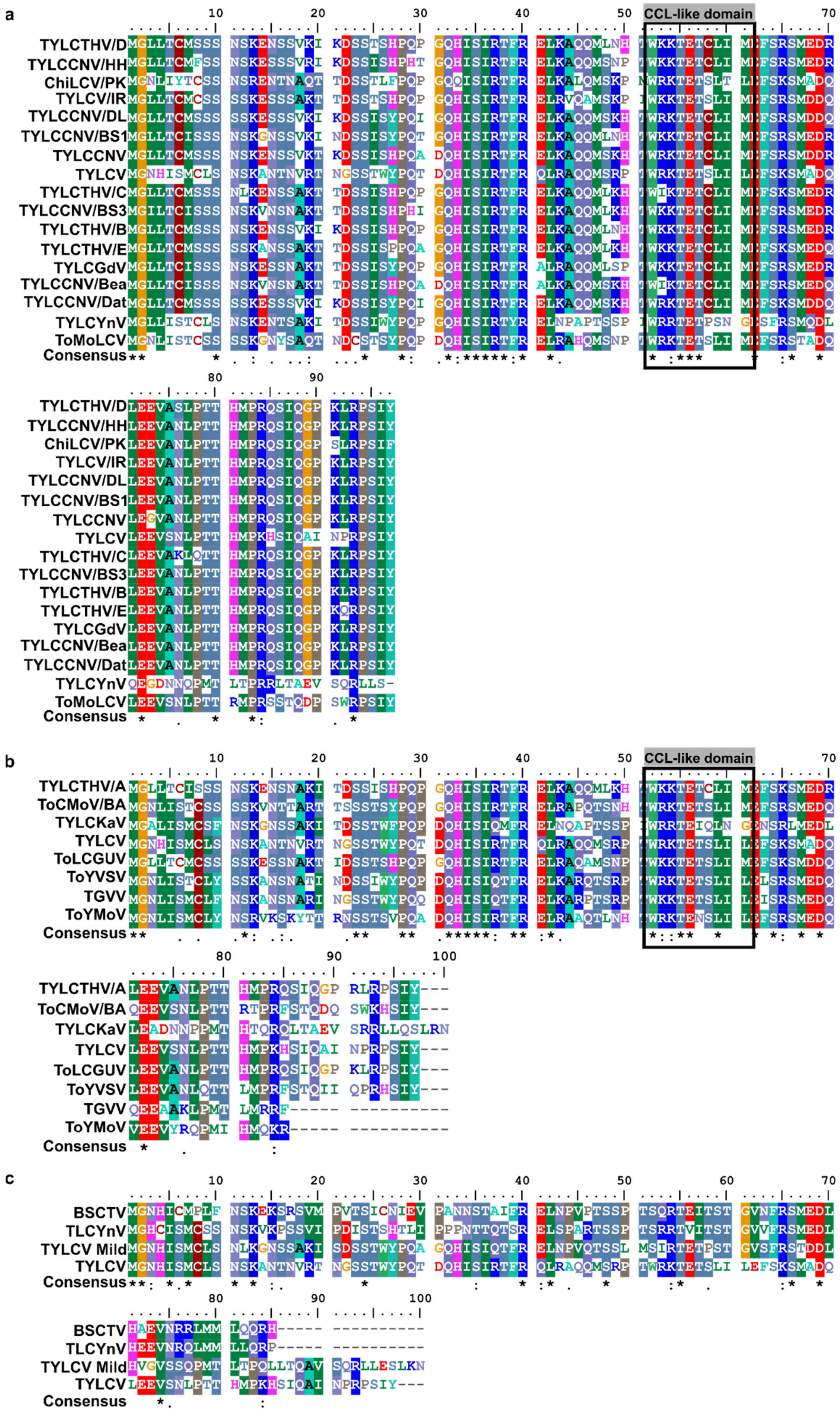


**Figure S4. The CCL-like domain is conserved in the C4 proteins from begomoviruses.**

a, b. Alignments between the C4 protein sequence from TYLCV and that from 16 selected monopartite begomoviruses causing tomato leaf curl disease (a), or that from 7 selected bipartite begomoviruses (b). In (b), TYLCV is shown for reference.

c. Alignment between C4 protein sequence from BSCTV (belonging to the *Curtovirus* genus), TLCYnV (which interacts with NbSKη), TYLCV-Mild, and TYLCV.

Multiple sequence alignments were performed by BioEdit software. The CCL-like sequence is framed in a black rectangle. All sequences were downloaded from GenBank; details regarding virus species, isolate, and accession numbers can be found in Table S7.

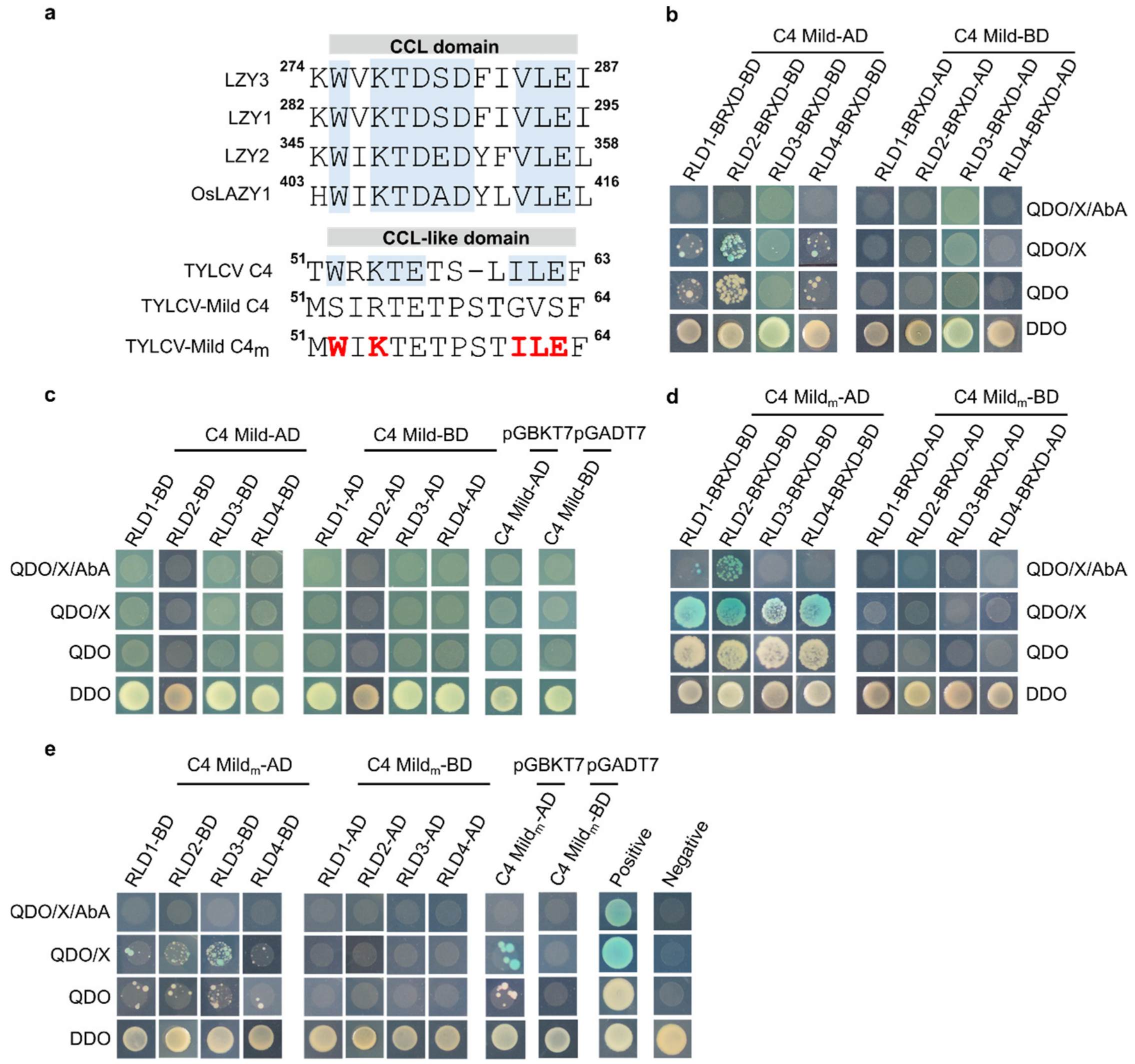


**Figure S5. The inclusion of a CCL-like domain in the CCL-like-free C4 from TYLCV-Mild is not sufficient to promote its interaction with RLDs.**

a. Design of C4 Mild$_m$. Key amino acids of the CCL-like domain of C4, highlighted in red bold letters, were introduced at equivalent positions in C4 Mild, generating C4 Mild$_m$, now containing a CCL-like domain. Numbers indicate amino acid positions in the corresponding proteins.

b. Interaction between C4 Mild and the BRX domain (BRXD) of Arabidopsis RLD1-4, tested by yeast two-hybrid (Y2H).

c. Interaction between C4 Mild and full-length Arabidopsis RLD1-4, tested by Y2H.

d. Interaction between C4 Mild$_m$ and the BRX domain (BRXD) of Arabidopsis RLD1-4, tested by Y2H.

e. Interaction between C4 Mild$_m$ and full-length Arabidopsis RLD1-4, tested by Y2H.

These experiments were performed three times with similar results; one representative replicate is shown here.

C4 Mild, C4 from TYLCV-Mild; C4, C4 from TYLCV.

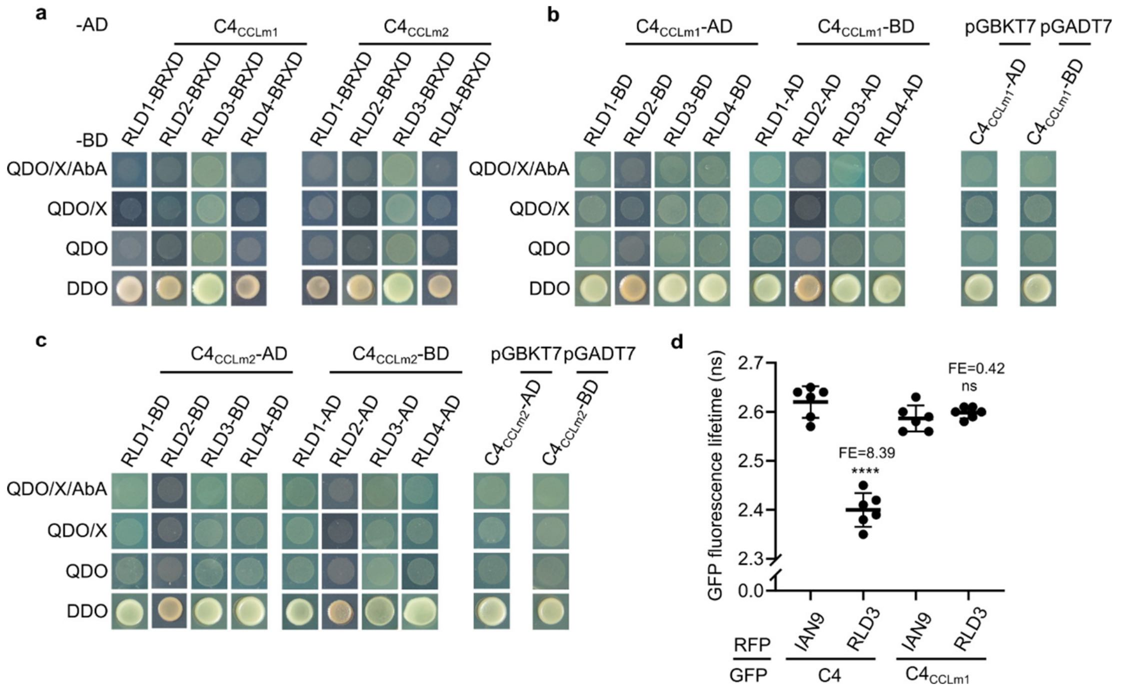


**Figure S6. $C4_{CCLm1}$ and $C4_{CCLm2}$ do not interact with RLD proteins.**

a. Interaction between $C4_{CCLm1}$/$C4_{CCLm2}$ and the BRX domain (BRXD) of Arabidopsis RLD1-4, tested by yeast two-hybrid (Y2H).

b. Interaction between $C4_{CCLm1}$ and full-length Arabidopsis RLD1-4, tested by Y2H.

c. Interaction between $C4_{CCLm2}$ and full-length RLD1-4, tested by Y2H.

d. Interaction between $C4_{CCLm1}$ and RLD3 upon transient co-expression in *N. benthamiana* leaves, analyzed by FRET-FLIM. FE, FRET efficiency. The membrane protein IAN9 is used as negative control. Samples were taken at 2 days post-agroinfiltration. Each dot represents the GFP fluorescence lifetime (ns, nanoseconds) of one technical replicate consisting of one field; the thick line indicates the average value, and the error bars correspond to standard deviations. Significant differences between groups were determined by one-way ANOVA ($P<0.0001$, F=80.63, df=3), followed by multiple comparisons of means by applying Tukey test; asterisks represent statistically significant differences at: ****, $P<0.0001$; ns, not significant.

All experiments in this figure were performed three times with similar results. One replicate is shown.

C4, C4 from TYLCV.

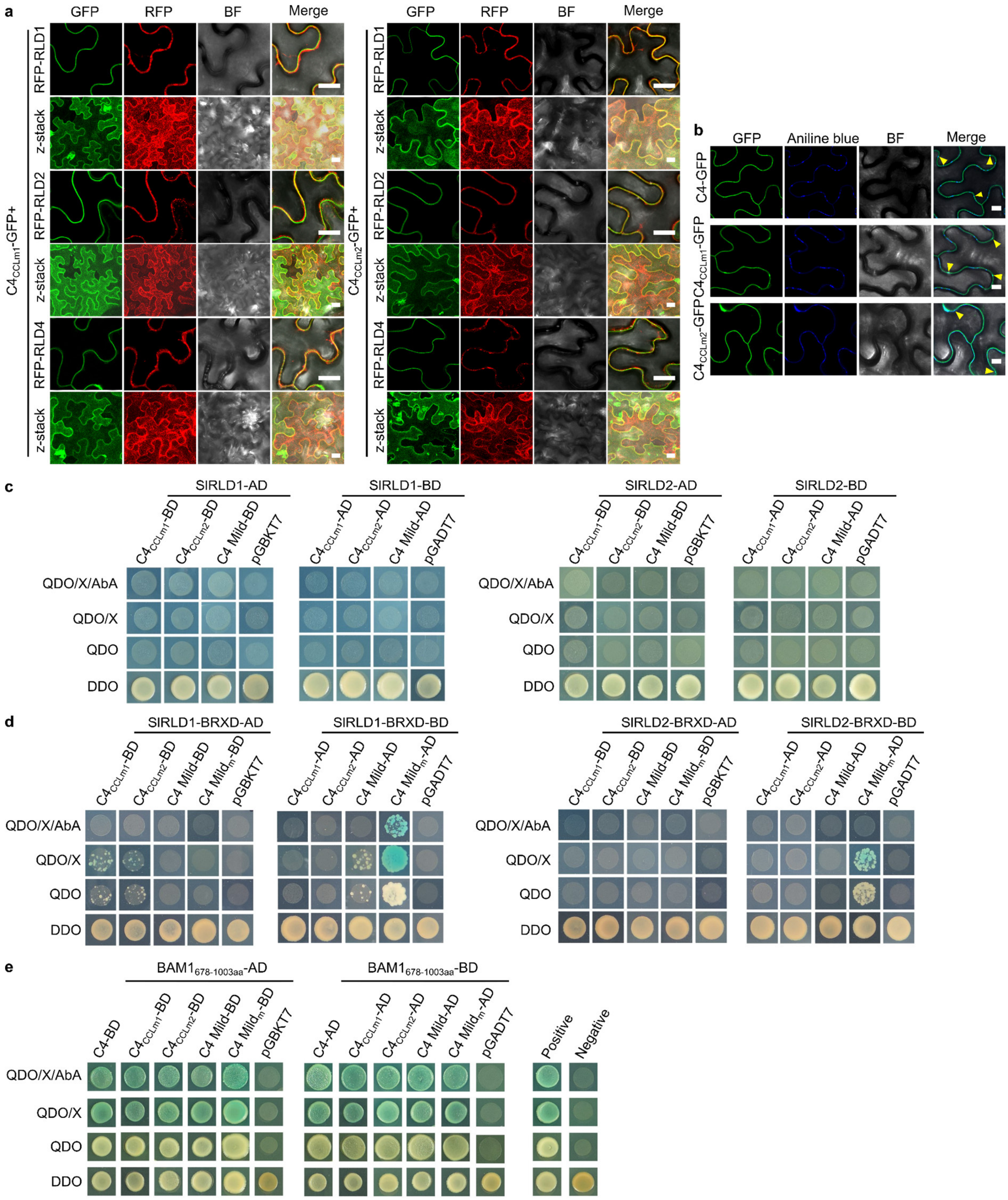


**Figure S7. An intact CCL-like domain is required for C4 from TYLCV to interact with RLDs, but dispensable for other interactions of C4 at the plasma membrane.**

a. Subcellular localization of Arabidopsis RFP-RLD1, 2, and 4 transiently expressed in *N. benthamiana* leaves in the presence or absence of $C4_{CCLm1}$/$C4_{CCLm2}$-GFP. White rectangles indicate close-ups.

b. Subcellular localization of C4-GFP/$C4_{CCLm1}$-GFP/$C4_{CCLm2}$-GFP upon transient expression in *N. benthamiana* leaves. Aniline blue staining labels callose deposits. Arrowheads indicate plasmodesmata.

In a,b, Images were taken at 2 days post-agroinfiltration. BF, bright field. Z-stack shows the maximum projection of a vertical cross-section through the observed cells. Scale bar, 20 μm.

c. Interaction between full-length tomato RLD1 and 2 (SlRLD1/2) and $C4_{CCLm1}$/$C4_{CCLm2}$/C4 Mild, tested by yeast two-hybrid (Y2H).

d. Interaction between the BRX domain (BRXD) of SlRLD1/SlRLD2 and $C4_{CCLm1}$/$C4_{CCLm2}$/C4 Mild/C4 $Mild_{m}$, tested by Y2H.

e. Interaction between the kinase domain (678-1003aa) of the plasma membrane-localized receptor kinase BAM1 and C4/$C4_{CCLm1}$/$C4_{CCLm2}$/C4 Mild/C4 $Mild_{m}$, tested by Y2H.

All experiments were performed three times with similar results; one representative replicate is shown here.

C4, C4 from TYLCV, C4 Mild, C4 Mild from TYLCV-Mild

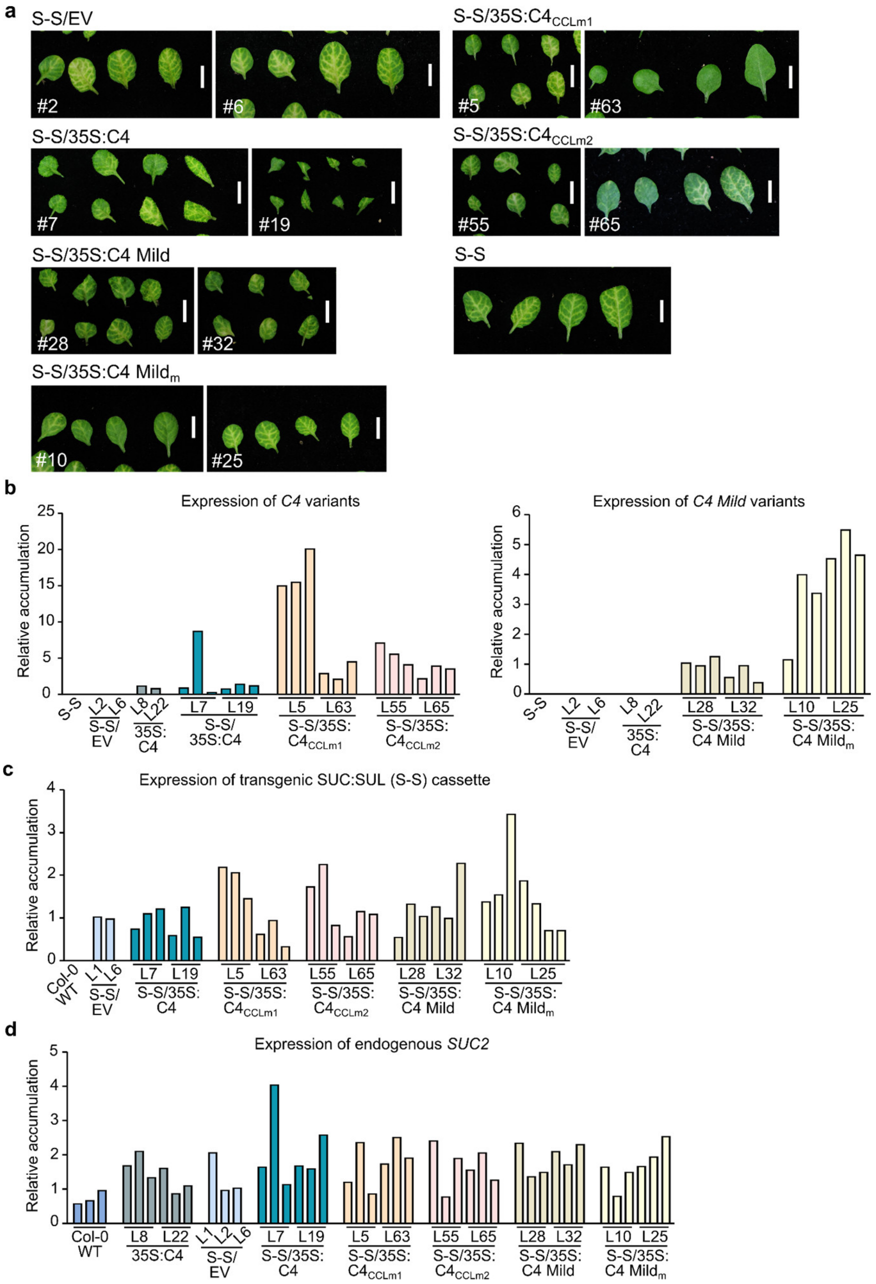


**Figure S8. Bleaching spread and transgene expression in SUC:SUL Arabidopsis plants transformed with C4 variants.**

a. Representative images of bleaching spread phenotype in leaves of 4-week-old SUC:SUL (S-S) plants expressing C4, C4 Mild, or their respective mutant forms (T2 generation). Scale bar, 1 cm. S-S/EV as well as non-transformed S-S leaves are shown as controls.

b. Expression levels of *C4* variants (WT and mutants) in the transgenic plants shown in (a), measured by RT-qPCR. Note that reference level (set to 1) for C4 variants is C4 WT (averaged L8+L22, T3), whereas for C4 Mild variants is C4 Mild WT (averaged L28, T3).

c. Expression level of the *SUL-ChSA* (chalcone synthase intron) construct from the transgenic SUC:SUL cassette, in the transgenic plants shown in (a), by RT-qPCR. Note that reference level (set to 1) corresponds to S-S/EV (average L1+L6).

d. Expression levels of the endogenous *SUL* gene in the transgenic plants shown in (a), measured by RT-qPCR. Note that reference level (set to 1) corresponds to WT.

In the graphs in b, c, d, each bar represents the value obtained per individual RNA sample, consisting of a pool of material coming from four different plants.

C4, C4 from TYLCV; C4 Mild, C4 from TYLCV-Mild; S-S, SUC:SUL plants; EV, empty vector.

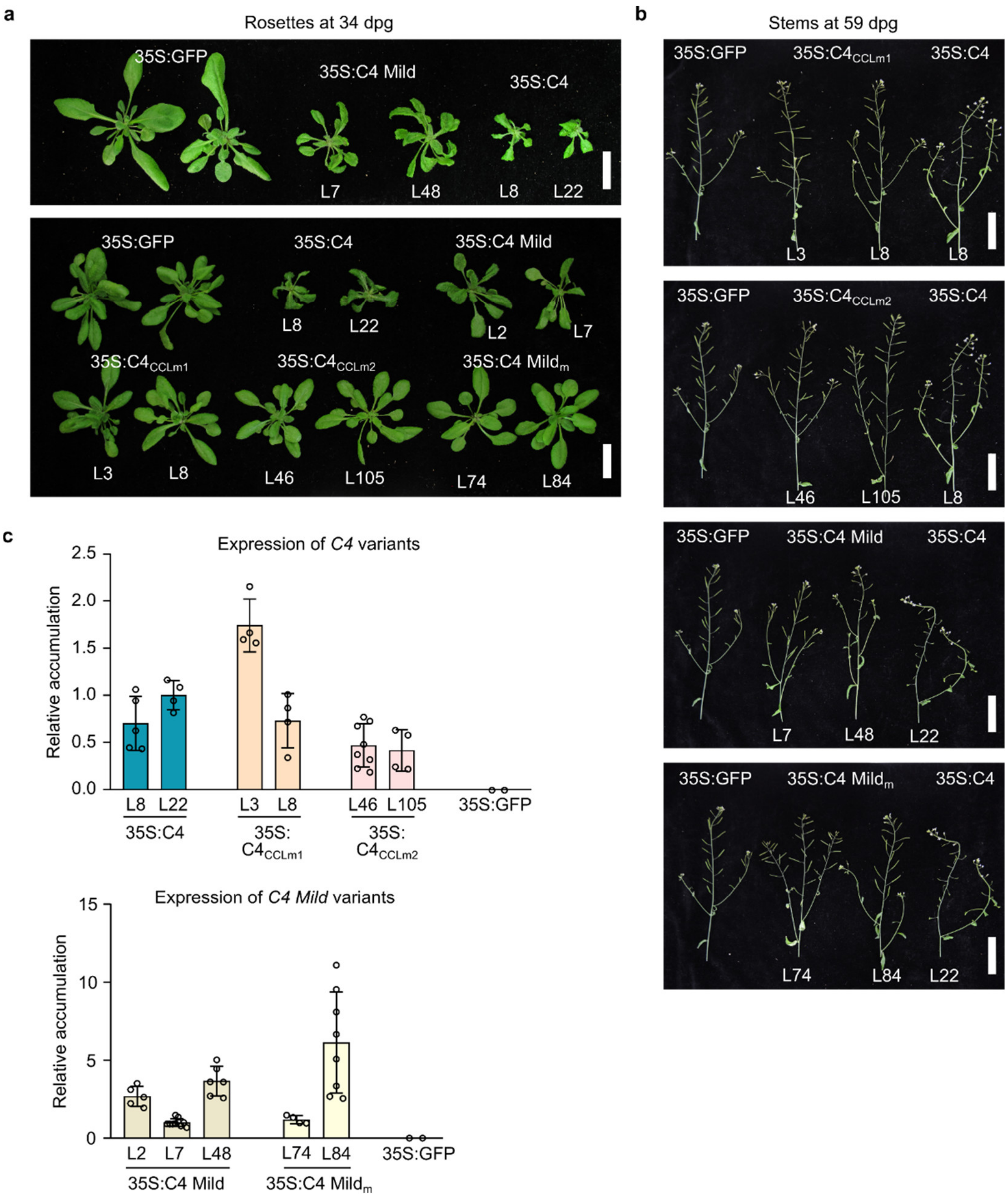


**Figure S9. Developmental phenotypes and transgene expression in Arabidopsis plants transformed with C4 variants.**

a, b. Representative phenotypes in rosettes of 34-days-old plants (a) and in stems, siliques, and inflorescences of 59-days-old plants (b) expressing C4, C4 Mild, or their respective mutant forms (T3 generation). Scale bar in (a), 2 cm; scale bar in (b), 4 cm.

c. Relative expression levels of the *C4* variants in the transgenic plants shown in (a), measured by RT-qPCR. Note that reference level (set to 1) for *C4* variants is 35S:C4 (L22, T3), whereas for *C4* Mild variants is 35S:C4 Mild (L7, T3). A stable line expressing GFP (35S:GFP) is shown as reference and as negative control in expression analyses. Note that the plants used in (b) as reference (35S:GFP as well as 35S:C4 lines) are the same in different pictures.

C4, C4 from TYLCV; C4 Mild, C4 from TYLCV-Mild; dpg, days post-germination.

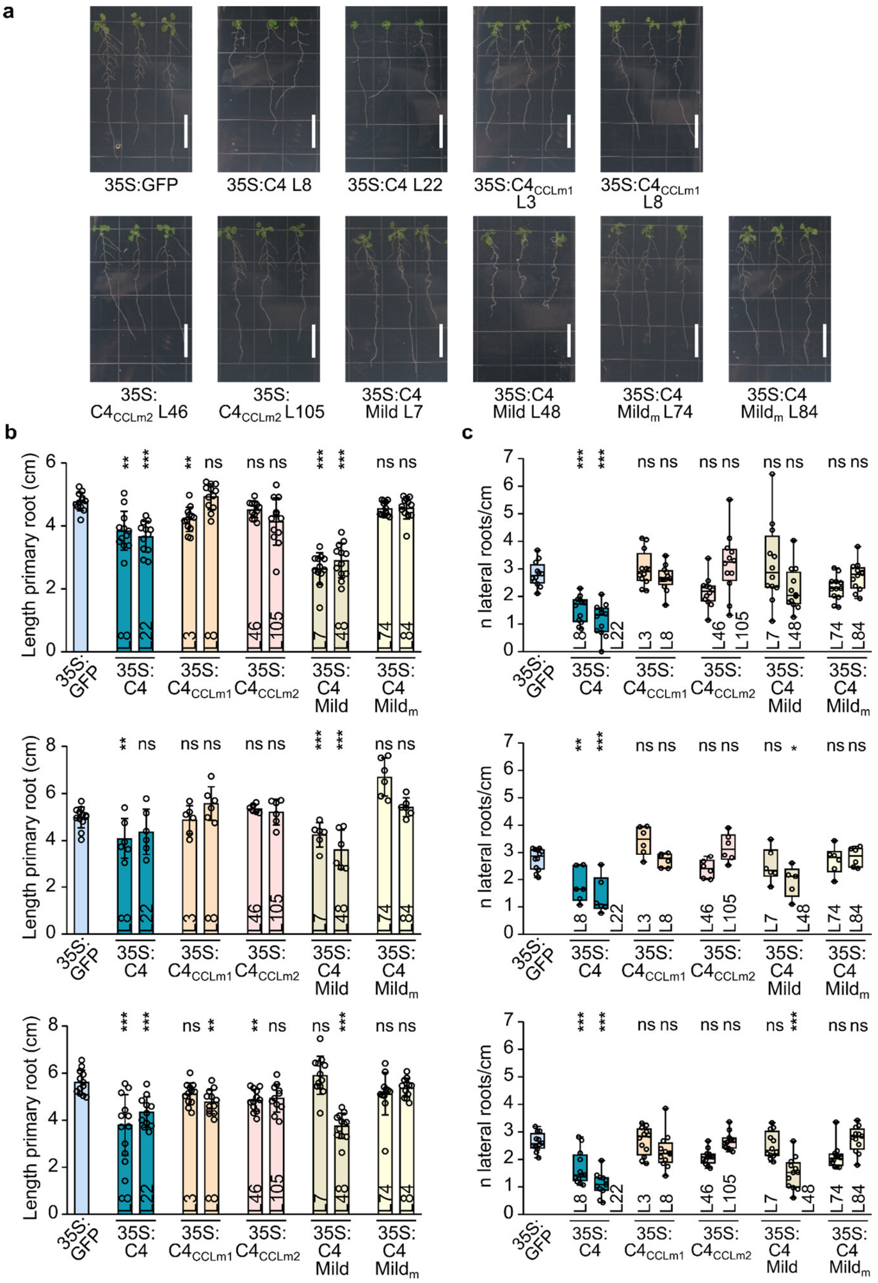


**Figure S10. Root phenotypes in Arabidopsis plants transformed with C4 variants.**

a. Representative root phenotypes in plants expressing C4, C4 Mild, or their respective mutant forms (T3 generation), at 14 days post-germination. Scale bar, 1 cm. A stable line expressing GFP (35S:GFP) is shown as reference. Note that some pictures in (a) are also included in Figure S12a as reference, since both experiments were run in parallel.

b, c. Quantifications of the primary root length (b) and of the number of lateral roots per unit of primary root length (c), as obtained in three independent experiments. In (b), each dot represents the root length of an individual plant, and the error bars indicate standard deviations; a minimum of 6 plants were analyzed per line. Significant differences between groups were determined by either one-way ANOVA (middle graph, *P*=2.19E-10, F=11.044, df=10) or Kruskal-Wallis (upper graph, *P*=2,029E-14, H=87.078, df=10; bottom graph, *P*=5.73E-10, H=64.213, df=10), followed by multiple comparisons of means between the C4-expressing lines against the reference group (35S:GFP), by applying either Dunnett's test (middle graph) or pairwise comparisons between each group and the reference group (upper and bottom graphs, Mann-Whitney U test, significance 0.005, after Bonferroni's correction for multiple comparisons); asterisks represent statistically significant differences at: ***, $P<0.001$; **, $P<0.01$; ns, not significant. In (c), each dot represents the number of lateral roots per unit of primary root length of an individual plant, and the error bars the lowest/highest values; a minimum of 6 plants were analyzed per line. Significant differences between groups were determined by one-way ANOVA (upper graph, $P$=5.70E-10, F=9.298, df=10; middle graph, $P$=1.14E-8, F=8.778, df=10; bottom graph, $P$=1.92E-17, F=15.487, df=10), followed by multiple comparisons of means between the C4-expressing lines against the reference group (35S:GFP), by applying Dunnett's test; asterisks represent statistically significant differences at: ***, $P<0.001$; **, $P<0.01$; ns, not significant.

C4, C4 from TYLCV; C4 Mild, C4 from TYLCV-Mild.

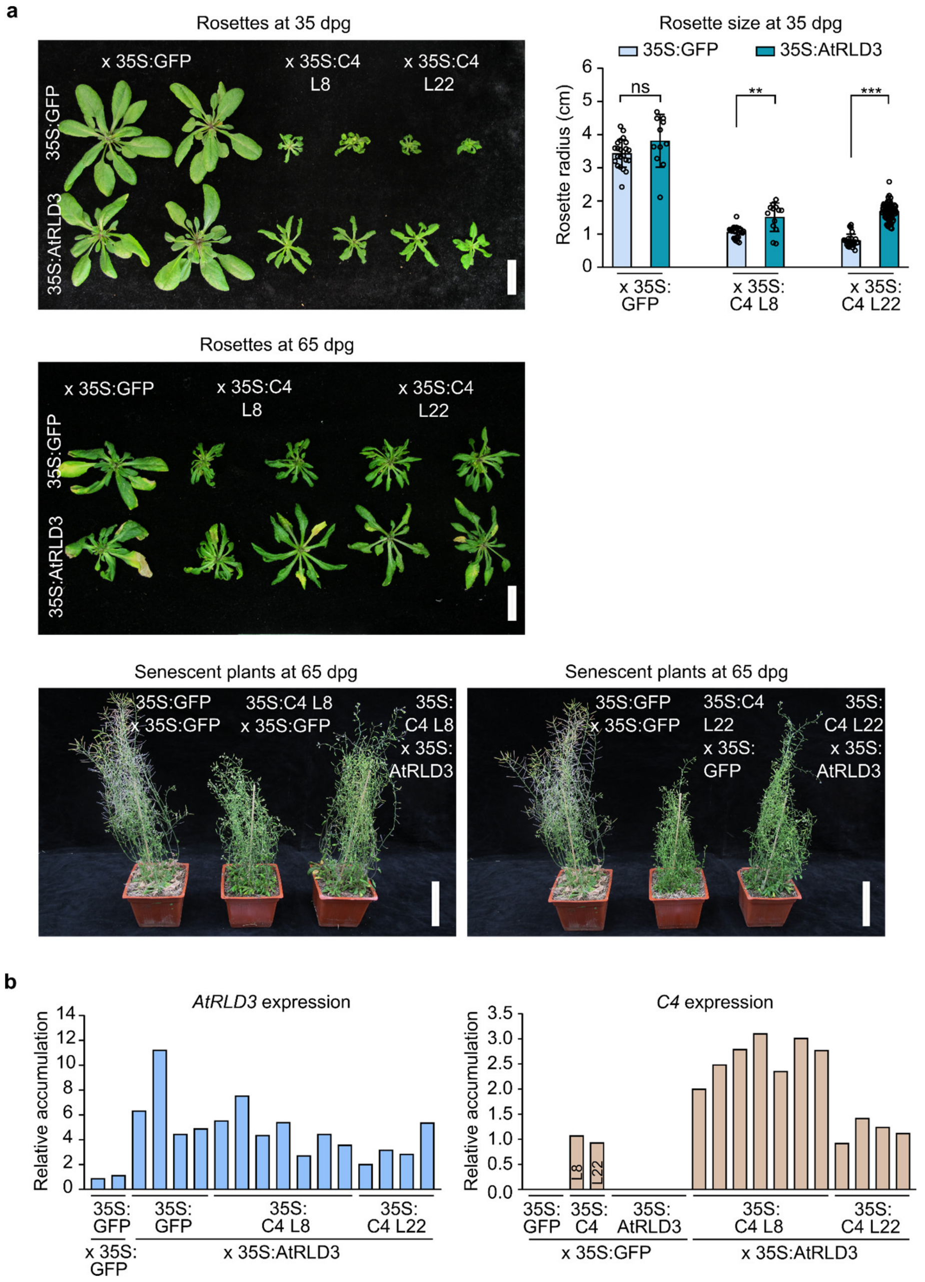


**Figure S11. Overexpression of *RLD3* alleviates C4-induced developmental phenotypes in Arabidopsis.**

a. Representative rosette phenotypes in the F1 generation plants resulting from crossing 35S:C4 lines 8 and 22 (T2) with 35S:AtRLD3 lines 20 and 10 (T2) parentals, respectively, at 35 days post-germination (dpg) (upper panel, on the left) or at 65 dpg (middle panel); scale bar, 2 cm. Complete plants (bottom panels) were also photographed at 65 dpg; scale bar, 5 cm. In these experiments, F1s coming from crossing a stable line expressing GFP (35S:GFP) with 35S:AtRLD3 T2 parentals or with itself are used as controls. The bar graph (upper panel, on the right) shows the quantification of the rosette average radius at 35 dpg; each dot represents the average of three radius measurements per rosette, and error bars indicate standard deviations. A minimum of 11 rosettes per genotype were analyzed, when possible. Differences between pairs of groups were assessed by applying Mann-Whitney U test; asterisks represent statistically significant differences at: ***, $P<0.001$; **, $P<0.01$; ns, not significant.

b. Relative expression levels of *AtRLD3* and *C4* in the plants shown in (a), measured by RT-qPCR. Note that reference level (set to 1) for *AtRLD3* is F1 35S:GFP x 35S:GFP, whereas for *C4* is 35S:C4 (averaged L8+22, T3).

C4, C4 from TYLCV; dpg, days post-germination.

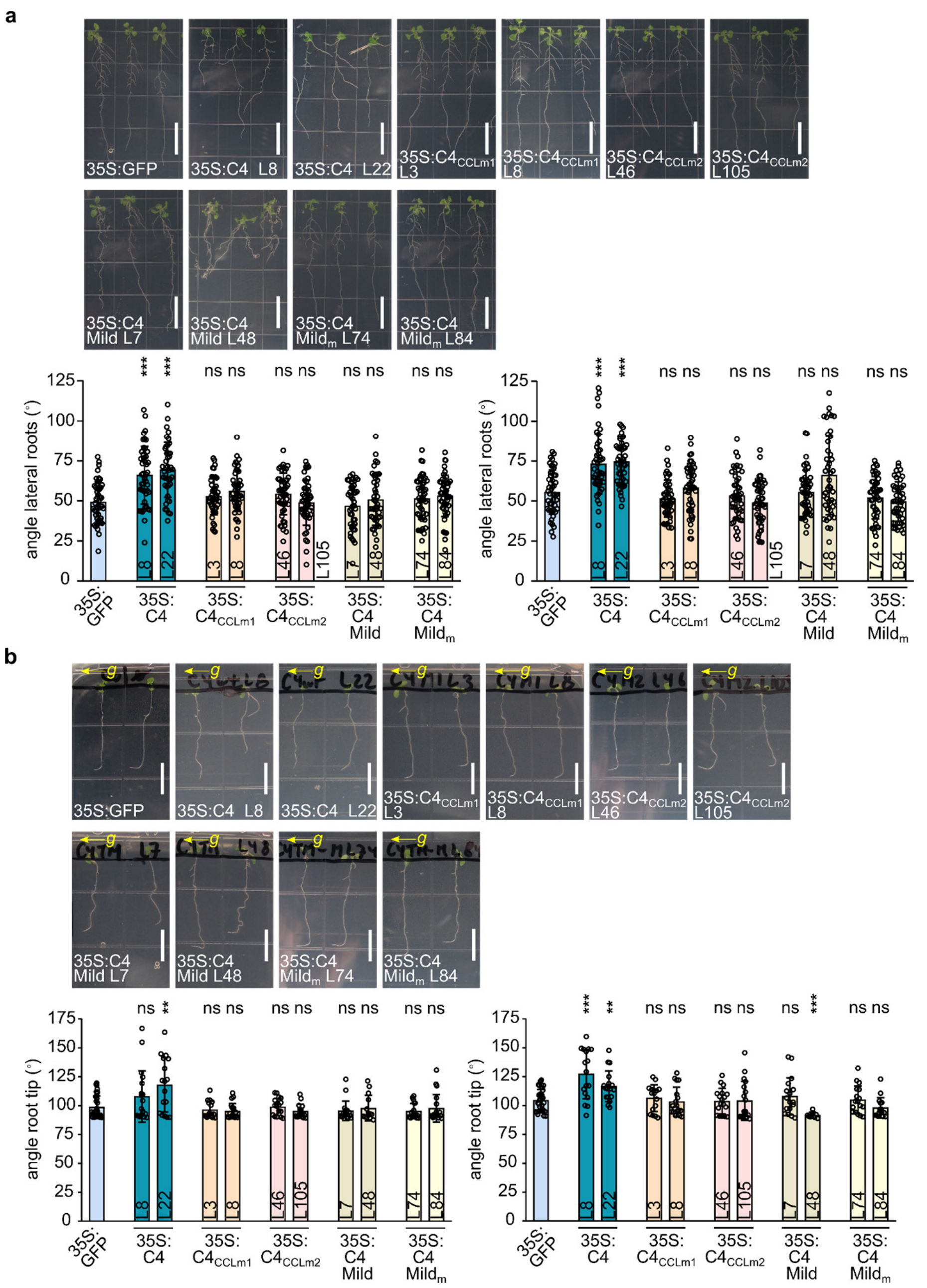


**Figure S12. Root gravitropic responses are altered in Arabidopsis plants expressing C4 from TYLCV.**

a. Pictures show representative root phenotypes in plants expressing C4, C4 Mild, or their respective mutant forms (T3 generation), at 14 days post-germination (dpg). Scale bar, 1 cm. A stable line expressing GFP (35S:GFP) is shown as reference. Graphs show quantifications of the lateral root angles relative to the main roots in the previous transgenic plants (gravitropic set-point angles, GSAs), in two independent experiments. Each dot represents the angle measurements of a lateral root, and error bars indicate standard deviations. Angles of four different roots per plant were measured, and a minimum of 6 plants were analyzed per line. Significant differences between groups were determined by one-way ANOVA (replicate on the left: *P*=2.77E-26, F=11.255, df=10; replicate on the right: *P*=1.86E-17, F=17.621, df=10), followed by multiple comparisons of means between the C4-expressing lines against the reference group (35S:GFP), by applying Dunnett's test; asterisks represent statistically significant differences at: ***, *P*<0.001; ns, not significant. Note that some pictures in (a) are also shown in Figure S10a, as reference, since both experiments were run in parallel.

b. Pictures show representative root tip angles in plants expressing C4, C4 Mild, or their respective mutant forms (T3 generation), at 7 dpg, after growing for 12 h rotated 90 degrees with respect to their initial growing position: the new *g* force direction is depicted in the pictures by the yellow arrow to the left. Scale bar, 1 cm. A stable line expressing GFP (35S:GFP) is shown as reference. Graphs show quantifications of the root tip angles in the previous transgenic plants, after receiving the gravitropic treatment, in two independent experiments. Each dot represents the individual measurement of a root tip angle, and error bars indicate standard deviations. Root tip angles of a minimum of 14 plants were analyzed per line. Differences between groups were assessed by applying Kruskal-Wallis test (replicate on the left: *P*=0.031, H=19.74, df=10; replicate on the right: *P*=3.046-07, H=49.67, df=10), followed by pairwise comparisons between each group and the reference group (35S:GFP) (Mann-Whitney U test, significance 0.005, after Bonferroni's correction for multiple comparisons); asterisks represent statistically significant differences at: ***, *P*<0.001; **, *P*=0.002; ns, not significant.

C4, C4 from TYLCV; C4 Mild, C4 from TYLCV-Mild.

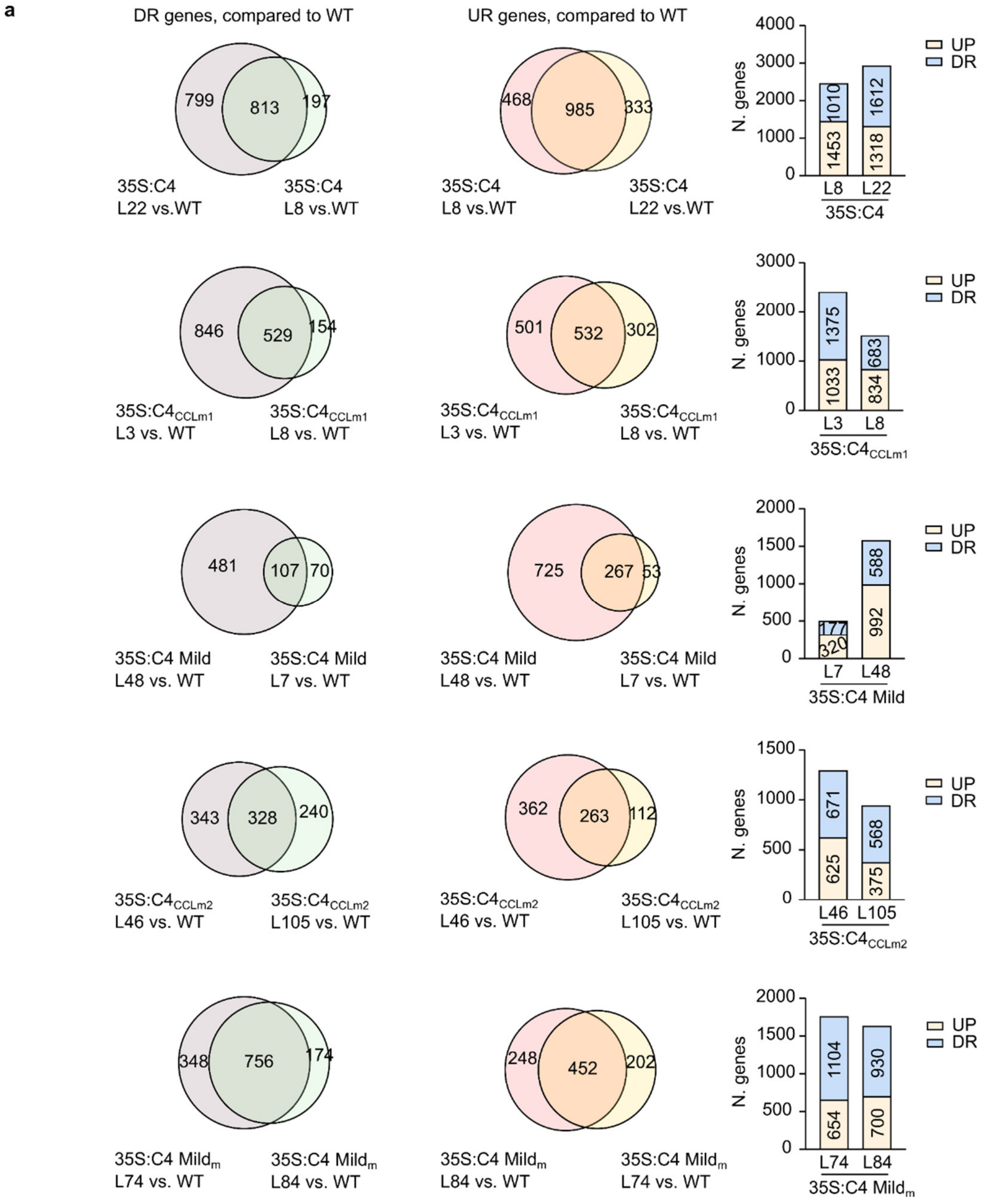

a
DR genes, compared to WT
UR genes, compared to WT
799
813
197
35S:C4
L22 vs.WT
35S:C4
L8 vs.WT
468
985
333
35S:C4
L8 vs.WT
35S:C4
L22 vs.WT
N. genes
UP
DR
1453
1010
1318
1612
L8
L22
35S:C4
846
529
154
35S:C4CCLm1
L3 vs. WT
35S:C4CCLm1
L8 vs. WT
501
532
302
35S:C4CCLm1
L3 vs. WT
35S:C4CCLm1
L8 vs. WT
1033
1375
834
683
L3
L8
35S:C4CCLm1
481
107
70
35S:C4 Mild
L48 vs. WT
35S:C4 Mild
L7 vs. WT
725
267
53
35S:C4 Mild
L48 vs. WT
35S:C4 Mild
L7 vs. WT
320
177
992
588
L7
L48
35S:C4 Mild
343
328
240
35S:C4CCLm2
L46 vs. WT
35S:C4CCLm2
L105 vs. WT
362
263
112
35S:C4CCLm2
L46 vs. WT
35S:C4CCLm2
L105 vs. WT
625
671
375
568
L46
L105
35S:C4CCLm2
348
756
174
35S:C4 Mildm
L74 vs. WT
35S:C4 Mildm
L84 vs. WT
248
452
202
35S:C4 Mildm
L84 vs. WT
35S:C4 Mildm
L74 vs. WT
654
1104
700
930
L74
L84
35S:C4 Mildm

**b**

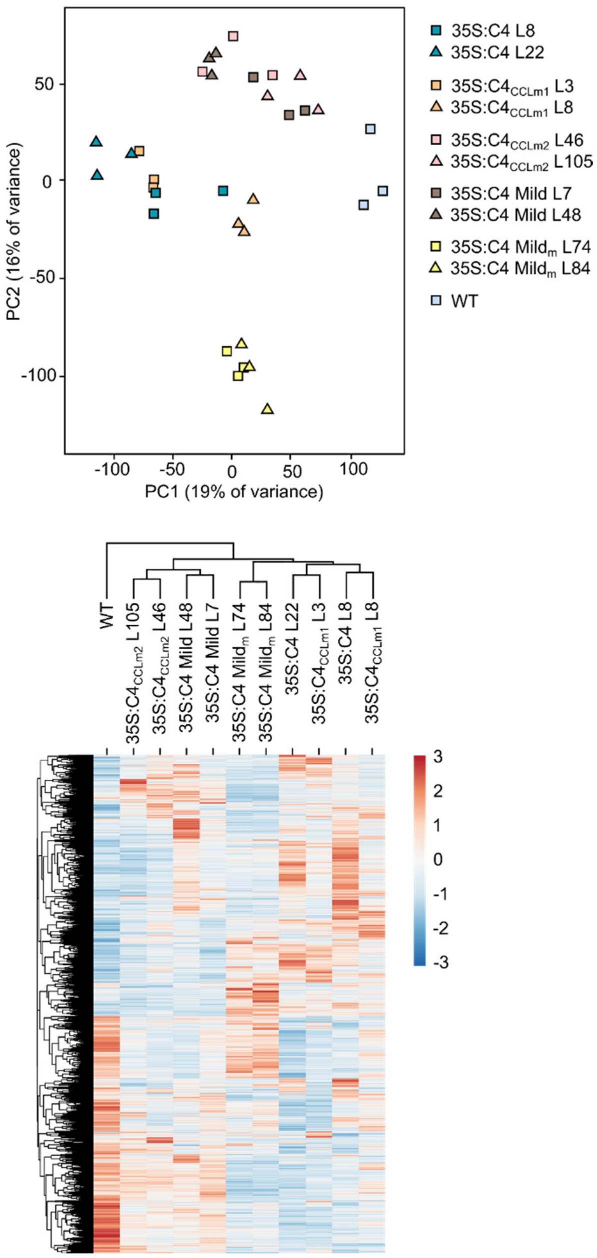

35S:C4 L8
35S:C4 L22
35S:C4$_{CCLm1}$ L3
35S:C4$_{CCLm1}$ L8
35S:C4$_{CCLm2}$ L46
35S:C4$_{CCLm2}$ L105
35S:C4 Mild L7
35S:C4 Mild L48
35S:C4 Mild$_m$ L74
35S:C4 Mild$_m$ L84
WT
PC1 (19% of variance)
PC2 (16% of variance)
50
0
-50
-100
-100
-50
0
50
100
WT
35S:C4$_{CCLm2}$ L105
35S:C4$_{CCLm2}$ L46
35S:C4 Mild L48
35S:C4 Mild L7
35S:C4 Mild$_m$ L74
35S:C4 Mild$_m$ L84
35S:C4 L22
35S:C4$_{CCLm1}$ L3
35S:C4 L8
35S:C4$_{CCLm1}$ L8
3
2
1
0
-1
-2
-3

c

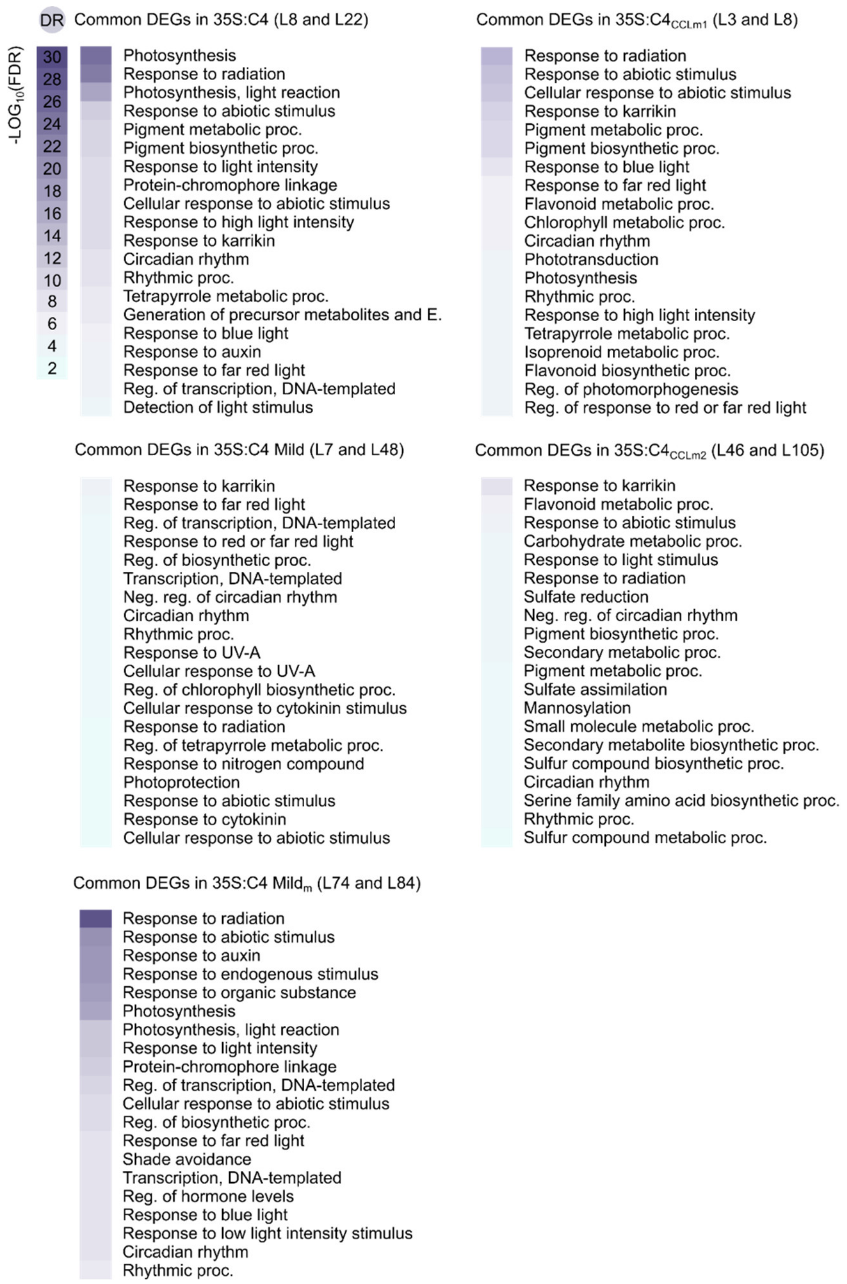
DR
-LOG10(FDR)
30
28
26
24
22
20
18
16
14
12
10
8
6
4
2
Common DEGs in 35S:C4 (L8 and L22)
Photosynthesis
Response to radiation
Photosynthesis, light reaction
Response to abiotic stimulus
Pigment metabolic proc.
Pigment biosynthetic proc.
Response to light intensity
Protein-chromophore linkage
Cellular response to abiotic stimulus
Response to high light intensity
Response to karrikin
Circadian rhythm
Rhythmic proc.
Tetrapyrrole metabolic proc.
Generation of precursor metabolites and E.
Response to blue light
Response to auxin
Response to far red light
Reg. of transcription, DNA-templated
Detection of light stimulus
Common DEGs in 35S:C4CCLm1 (L3 and L8)
Response to radiation
Response to abiotic stimulus
Cellular response to abiotic stimulus
Response to karrikin
Pigment metabolic proc.
Pigment biosynthetic proc.
Response to blue light
Response to far red light
Flavonoid metabolic proc.
Chlorophyll metabolic proc.
Circadian rhythm
Phototransduction
Photosynthesis
Rhythmic proc.
Response to high light intensity
Tetrapyrrole metabolic proc.
Isoprenoid metabolic proc.
Flavonoid biosynthetic proc.
Reg. of photomorphogenesis
Reg. of response to red or far red light
Common DEGs in 35S:C4 Mild (L7 and L48)
Response to karrikin
Response to far red light
Reg. of transcription, DNA-templated
Response to red or far red light
Reg. of biosynthetic proc.
Transcription, DNA-templated
Neg. reg. of circadian rhythm
Circadian rhythm
Rhythmic proc.
Response to UV-A
Cellular response to UV-A
Reg. of chlorophyll biosynthetic proc.
Cellular response to cytokinin stimulus
Response to radiation
Reg. of tetrapyrrole metabolic proc.
Response to nitrogen compound
Photoprotection
Response to abiotic stimulus
Response to cytokinin
Cellular response to abiotic stimulus
Common DEGs in 35S:C4CCLm2 (L46 and L105)
Response to karrikin
Flavonoid metabolic proc.
Response to abiotic stimulus
Carbohydrate metabolic proc.
Response to light stimulus
Response to radiation
Sulfate reduction
Neg. reg. of circadian rhythm
Pigment biosynthetic proc.
Secondary metabolic proc.
Pigment metabolic proc.
Sulfate assimilation
Mannosylation
Small molecule metabolic proc.
Secondary metabolite biosynthetic proc.
Sulfur compound biosynthetic proc.
Circadian rhythm
Serine family amino acid biosynthetic proc.
Rhythmic proc.
Sulfur compound metabolic proc.
Common DEGs in 35S:C4 Mildm (L74 and L84)
Response to radiation
Response to abiotic stimulus
Response to auxin
Response to endogenous stimulus
Response to organic substance
Photosynthesis
Photosynthesis, light reaction
Response to light intensity
Protein-chromophore linkage
Reg. of transcription, DNA-templated
Cellular response to abiotic stimulus
Reg. of biosynthetic proc.
Response to far red light
Shade avoidance
Transcription, DNA-templated
Reg. of hormone levels
Response to blue light
Response to low light intensity stimulus
Circadian rhythm
Rhythmic proc.

d

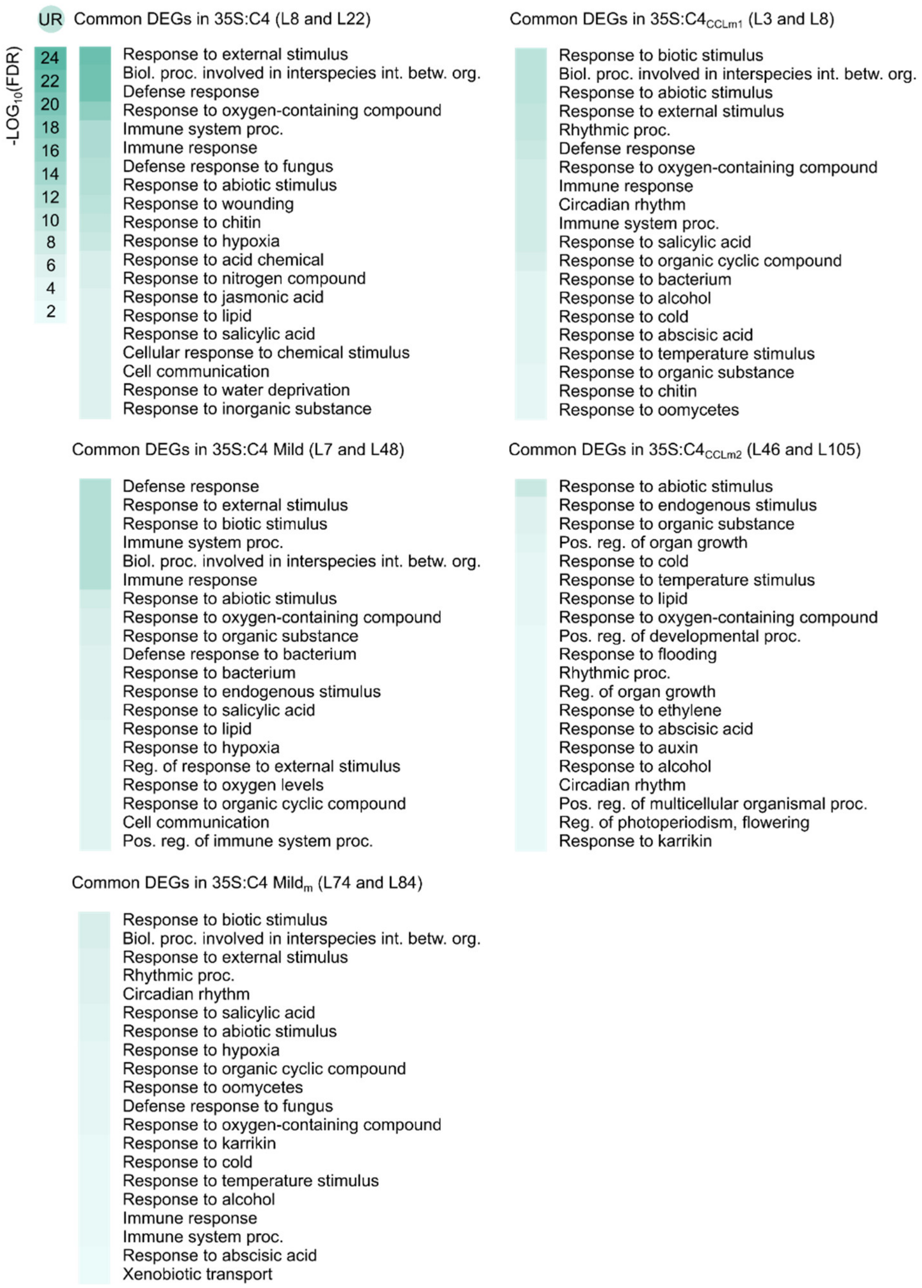
UR
-LOG10(FDR)
24
22
20
18
16
14
12
10
8
6
4
2
Common DEGs in 35S:C4 (L8 and L22)
Response to external stimulus
Biol. proc. involved in interspecies int. betw. org.
Defense response
Response to oxygen-containing compound
Immune system proc.
Immune response
Defense response to fungus
Response to abiotic stimulus
Response to wounding
Response to chitin
Response to hypoxia
Response to acid chemical
Response to nitrogen compound
Response to jasmonic acid
Response to lipid
Response to salicylic acid
Cellular response to chemical stimulus
Cell communication
Response to water deprivation
Response to inorganic substance
Common DEGs in 35S:C4CCLm1 (L3 and L8)
Response to biotic stimulus
Biol. proc. involved in interspecies int. betw. org.
Response to abiotic stimulus
Response to external stimulus
Rhythmic proc.
Defense response
Response to oxygen-containing compound
Immune response
Circadian rhythm
Immune system proc.
Response to salicylic acid
Response to organic cyclic compound
Response to bacterium
Response to alcohol
Response to cold
Response to abscisic acid
Response to temperature stimulus
Response to organic substance
Response to chitin
Response to oomycetes
Common DEGs in 35S:C4 Mild (L7 and L48)
Defense response
Response to external stimulus
Response to biotic stimulus
Immune system proc.
Biol. proc. involved in interspecies int. betw. org.
Immune response
Response to abiotic stimulus
Response to oxygen-containing compound
Response to organic substance
Defense response to bacterium
Response to bacterium
Response to endogenous stimulus
Response to salicylic acid
Response to lipid
Response to hypoxia
Reg. of response to external stimulus
Response to oxygen levels
Response to organic cyclic compound
Cell communication
Pos. reg. of immune system proc.
Common DEGs in 35S:C4CCLm2 (L46 and L105)
Response to abiotic stimulus
Response to endogenous stimulus
Response to organic substance
Pos. reg. of organ growth
Response to cold
Response to temperature stimulus
Response to lipid
Response to oxygen-containing compound
Pos. reg. of developmental proc.
Response to flooding
Rhythmic proc.
Reg. of organ growth
Response to ethylene
Response to abscisic acid
Response to auxin
Response to alcohol
Circadian rhythm
Pos. reg. of multicellular organismal proc.
Reg. of photoperiodism, flowering
Response to karrikin
Common DEGs in 35S:C4 Mildm (L74 and L84)
Response to biotic stimulus
Biol. proc. involved in interspecies int. betw. org.
Response to external stimulus
Rhythmic proc.
Circadian rhythm
Response to salicylic acid
Response to abiotic stimulus
Response to hypoxia
Response to organic cyclic compound
Response to oomycetes
Defense response to fungus
Response to oxygen-containing compound
Response to karrikin
Response to cold
Response to temperature stimulus
Response to alcohol
Immune response
Immune system proc.
Response to abscisic acid
Xenobiotic transport

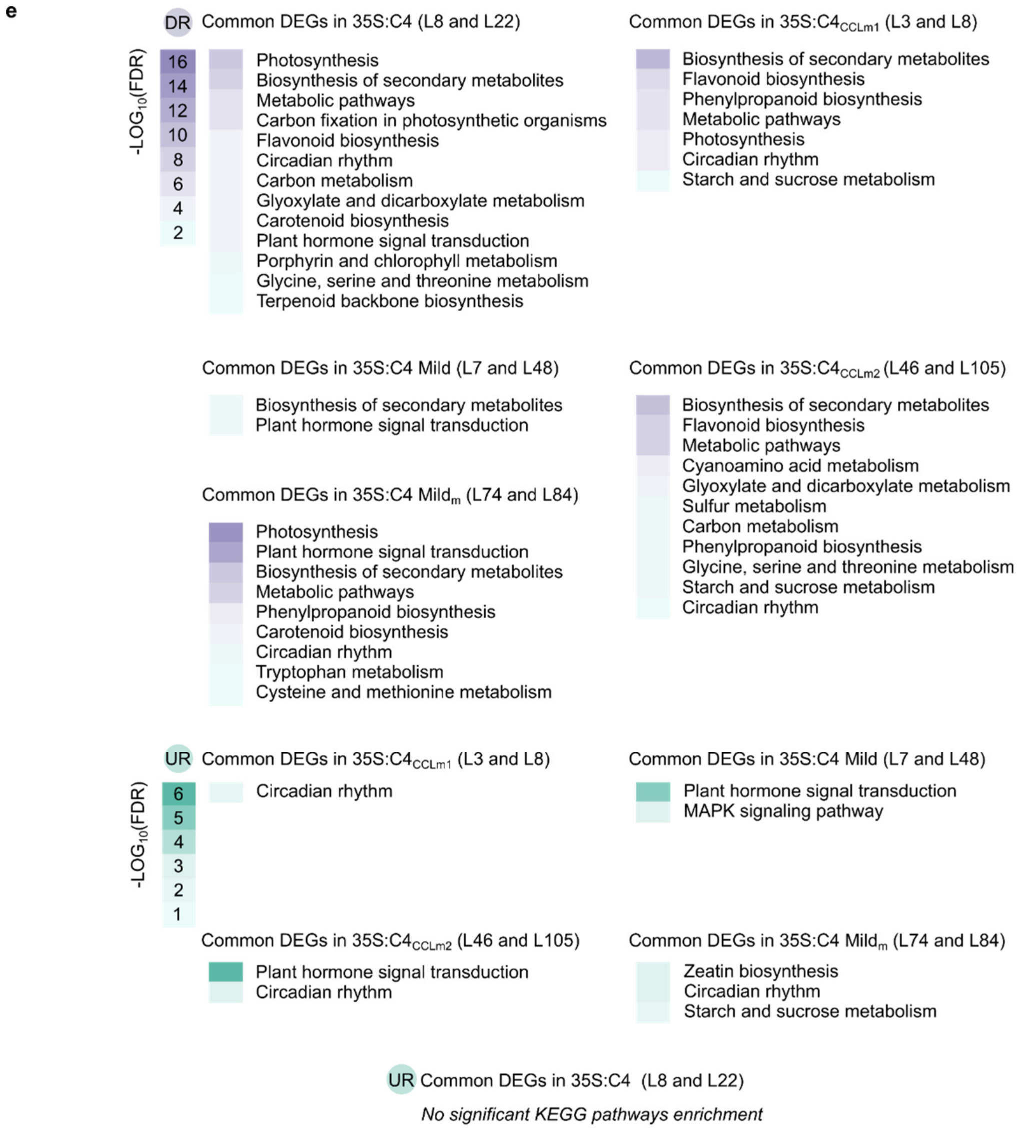
e
DR Common DEGs in 35S:C4 (L8 and L22)
-LOG10(FDR)
16
14
12
10
8
6
4
2
Photosynthesis
Biosynthesis of secondary metabolites
Metabolic pathways
Carbon fixation in photosynthetic organisms
Flavonoid biosynthesis
Circadian rhythm
Carbon metabolism
Glyoxylate and dicarboxylate metabolism
Carotenoid biosynthesis
Plant hormone signal transduction
Porphyrin and chlorophyll metabolism
Glycine, serine and threonine metabolism
Terpenoid backbone biosynthesis
Common DEGs in 35S:C4CCLm1 (L3 and L8)
Biosynthesis of secondary metabolites
Flavonoid biosynthesis
Phenylpropanoid biosynthesis
Metabolic pathways
Photosynthesis
Circadian rhythm
Starch and sucrose metabolism
Common DEGs in 35S:C4 Mild (L7 and L48)
Biosynthesis of secondary metabolites
Plant hormone signal transduction
Common DEGs in 35S:C4CCLm2 (L46 and L105)
Biosynthesis of secondary metabolites
Flavonoid biosynthesis
Metabolic pathways
Cyanoamino acid metabolism
Glyoxylate and dicarboxylate metabolism
Sulfur metabolism
Carbon metabolism
Phenylpropanoid biosynthesis
Glycine, serine and threonine metabolism
Starch and sucrose metabolism
Circadian rhythm
Common DEGs in 35S:C4 Mildm (L74 and L84)
Photosynthesis
Plant hormone signal transduction
Biosynthesis of secondary metabolites
Metabolic pathways
Phenylpropanoid biosynthesis
Carotenoid biosynthesis
Circadian rhythm
Tryptophan metabolism
Cysteine and methionine metabolism
UR Common DEGs in 35S:C4CCLm1 (L3 and L8)
-LOG10(FDR)
6
5
4
3
2
1
Circadian rhythm
Common DEGs in 35S:C4 Mild (L7 and L48)
Plant hormone signal transduction
MAPK signaling pathway
Common DEGs in 35S:C4CCLm2 (L46 and L105)
Plant hormone signal transduction
Circadian rhythm
Common DEGs in 35S:C4 Mildm (L74 and L84)
Zeatin biosynthesis
Circadian rhythm
Starch and sucrose metabolism
UR Common DEGs in 35S:C4 (L8 and L22)
No significant KEGG pathways enrichment

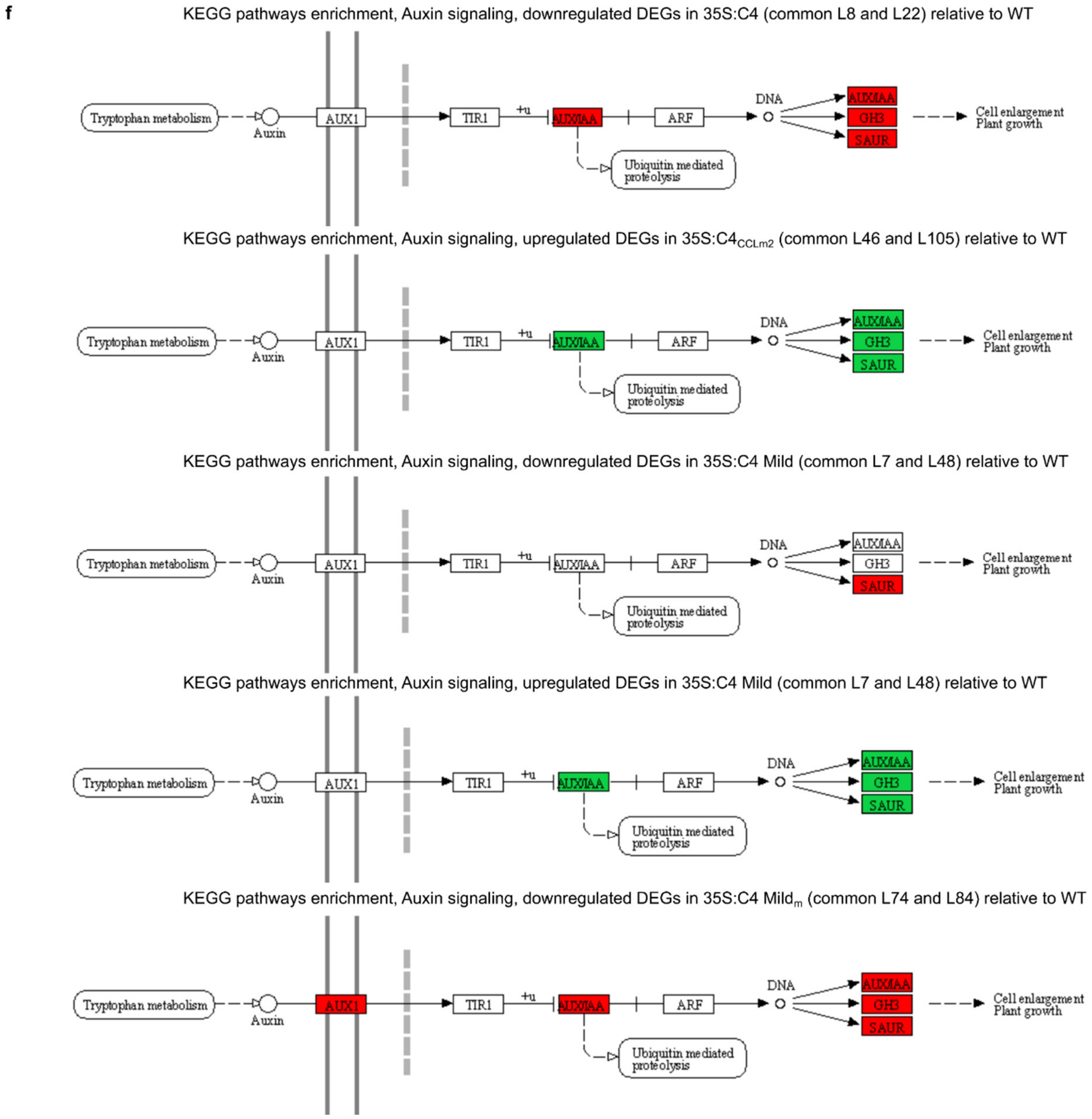


**Figure S13. Transcriptomic analyses of C4-expressing Arabidopsis plants.** Transcriptomic analyses were performed in aerial parts of 12-days-old Arabidopsis plants expressing C4 from TYLCV (C4), C4 from TYLCV-Mild (C4 Mild), or their respective mutant forms (T3 generation). Three biological replicates, each of them including material of at least 30 plants, were subjected to massive RNA sequencing, and comparative analyses of total RNAseq data were performed between each mutant line and the reference line (Col-0 WT plants). The summary of the total RNAseq data can be found in Table S2; counts and RPKM mapping to the C4 transgene can be found in Table S3.

a. Venn diagrams show the total number of genes found as differentially expressed (DEGs, fold-change=2, FDR<0.05), either downregulated (Venn diagrams on the left) or upregulated (Venn diagrams on the right) per transgenic line analyzed, as well as the numbers of overlapping genes found as differentially expressed in both transgenic lines. On the right, the stacked bar graphs summarize the total number of DEGs, either down- or upregulated, per line analyzed. Complete lists of DEGs can be found in Table S4.

b. Upper panel shows the principal component analysis (PC) graph, with PC1 and PC2 explaining 19% and 16%, respectively, of the total variance in the transcriptome data; bottom panel shows heatmap and hierarchical clustering of samples in (a); colour scales indicate Z-scores.

c, d. Functional enrichment analyses of downregulated (c) or upregulated (d) DEGs, in the indicated samples: the lists show the first 20 significantly overrepresented Gene Ontology (GO) categories (FDR<0.05) from the Biological Process ontology, after discarding low-level, redundant categories; colour codes and scales represent -$LOG_{10}$FDR values. For a full list of raw GO categories per genotype, see Table S5.

e. Enrichment analyses of DEGs (down- or upregulated genes) in The Kyoto Encyclopaedia of Genes and Genomes (KEGG) pathways: the lists show significantly overrepresented KEGG categories (FDR<0.05). Colour codes and scales represent -$LOG_{10}$FDR values. Full lists of genes and KEGG categories per genotype can be checked in Table S6. Note that no significant KEGG pathways enrichment was found for the upregulated DEGs in 35S:C4 genotype.

f, Visualization through Pathview of the 'Plant hormone signal transduction' KEGG pathway relative to auxin signaling, when found significantly enriched (FDR<0.05), for downregulated (35S:C4, 35S:C4 Mild and 35S:C4 $Mild_m$ genotypes, highlighted in red color) or upregulated (35S:C4 Mild and 35S:$C4_{CCLm2}$, highlighted in green color) DEGs.

DEGs, differentially expressed genes (here, when compared to WT); DR, downregulated DEGs; UP, upregulated DEGs; C4, C4 from TYLCV; C4 Mild, C4 from TYLCV-Mild; WT, Col-0 WT. Common abbreviations used in GO categories: E., Energy; Reg., Regulation; proc., process; Neg., Negative; Biol., Biological; int., interactions; betw., between; org., organisms; Pos., Positive.

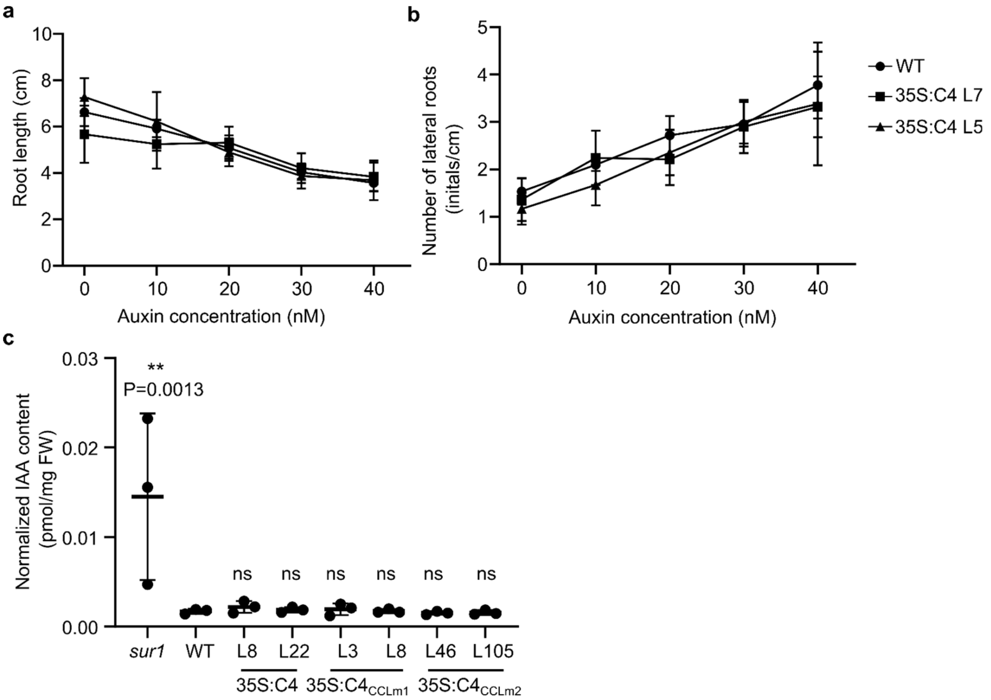


**Figure S14. C4 from TYLCV does not affect responses to exogenously applied IAA nor accumulation of IAA in transgenic Arabidopsis plants.**

a, b. C4-expressing plants (T3 generation) were grown in vertical plates supplemented with increasing concentrations of IAA (0-40 nM) and, at 12 days post-germination, primary root length (a) as well as the number of lateral roots per cm of primary root (b) were determined by Image J. In the graph, each dot represents the average, and the error bars the standard deviations; a minimum of ten roots were analyzed per line, and the experiment was repeated two times with similar results.

c. IAA content in 11-day-old whole seedlings of transgenic plants expressing C4 or its mutant forms $C4_{CCLm1}$ and $C4_{CCLm2}$. As controls, WT plants as well as the IAA over-accumulating mutant *sur1* were included. In the graph, each dot represents the IAA content of one biological replicate. The horizontal line represents the average IAA content, and the error bars the standard deviations. Significant differences between groups were determined by one-way ANOVA ($P$=0.0022, F=5.542, df=7), followed by multiple comparisons of means between the C4-expressing lines and the over-accumulating line *sur1* against the reference group (WT), by applying Dunnett's test; asterisks represent statistically significant differences at: **, $P$<0.01; ns, not significant.

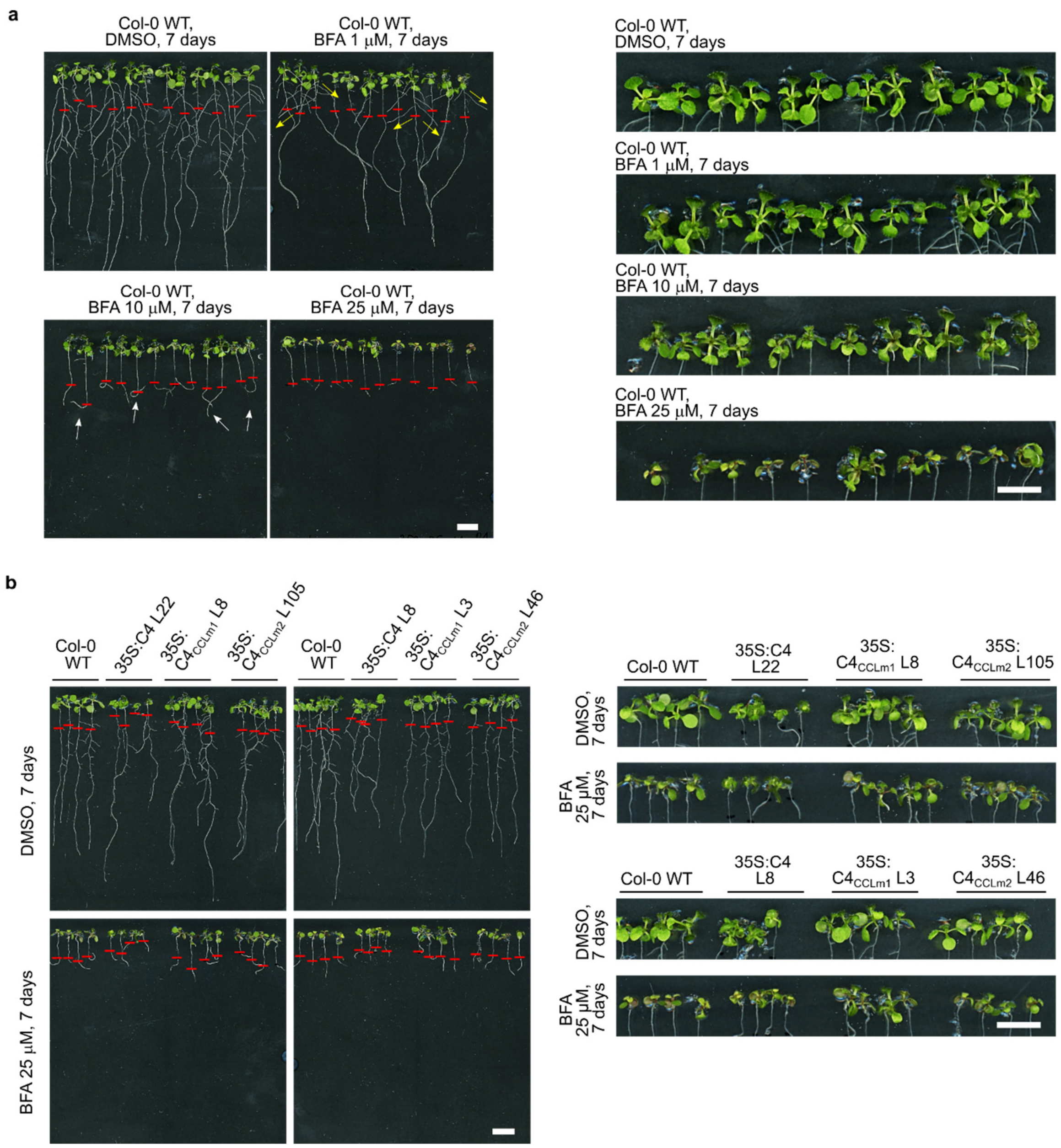


**Figure S15. BFA (Brefeldin A)-treated Arabidopsis seedlings show developmental alterations reminiscent of those induced by transgenic expression of C4 from TYLCV.**

a. 5-day-old wild-type (WT) Arabidopsis plants were transferred to ½ MS vertical plates supplemented with different concentrations of BFA (1, 10, or 25 μM dissolved in DMSO) or DMSO (control) for 7 days; pictures were taken at 12 days post-germination (dpg). On the left, pictures show root growth; note that BFA treatment induced the appearance of C4-like root phenotypes, characterized by wider lateral root angles (depicted with yellow arrows at 1 μM), aberrant root tip angles (depicted with white arrows at 10 μM), or curly rosette leaves (detailed pictures on the right).

b. Similar experiments as the ones described above were carried out with Arabidopsis plants expressing C4 or its mutant forms $C4_{CCLm1}$ and $C4_{CCLm2}$ (T3 generation), in the presence of 25 μM BFA. Representative phenotypes of roots (on the left) and rosettes (on the right) are presented. Red lines in both (a) and (b) mark the root size before BFA treatment, as reference. Scale bar, 1 cm. BFA, Brefeldin A; C4, C4 from TYLCV.

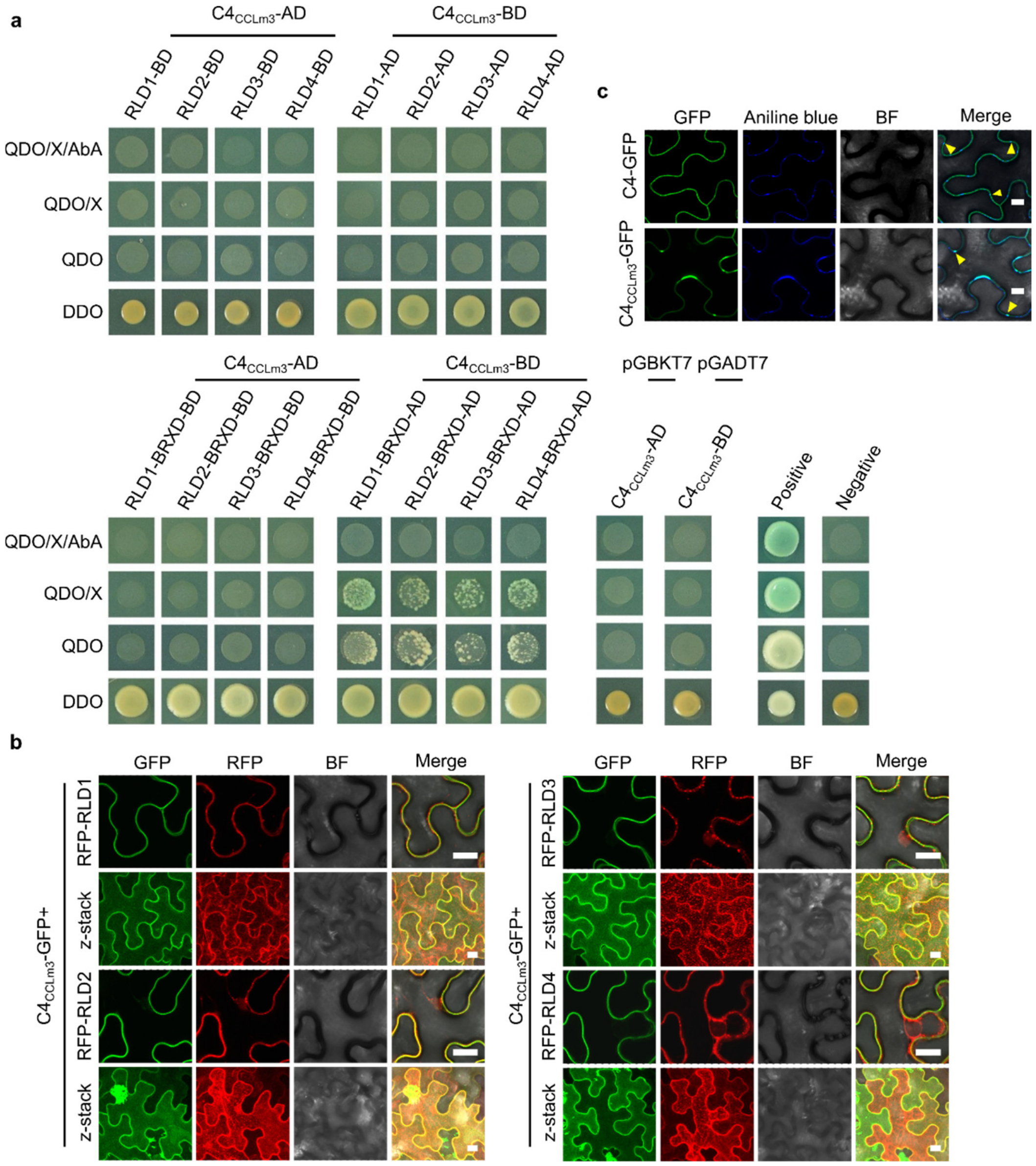


**Figure S16. C4$_{CCLm3}$ does not interact with RLD proteins nor recruit them to the plasma membrane.**

a. Interaction between C4$_{CCLm3}$ and full-length Arabidopsis RLD1-4 or their BRX domain (BRXD), tested by yeast two-hybrid (Y2H).

b. Subcellular localization of Arabidopsis RFP-RLD1-4 transiently expressed in *N. benthamiana* leaves in the presence or absence of C4$_{CCLm3}$-GFP. Images were taken at 2 days post-agroinfiltration (dpa).

c. Subcellular localization of C4-GFP/C4$_{CCLm3}$-GFP upon transient expression in *N. benthamiana* leaves. Aniline blue labels callose deposits. Arrowheads indicate plasmodesmata. Images were taken at 2 dpa.

BF, bright field. Z-stack shows the maximum projection of a vertical cross-section through the observed cells. Scale bar, 20 µm. All experiments in this figure were performed three time with similar results; one representative replicate is shown here.

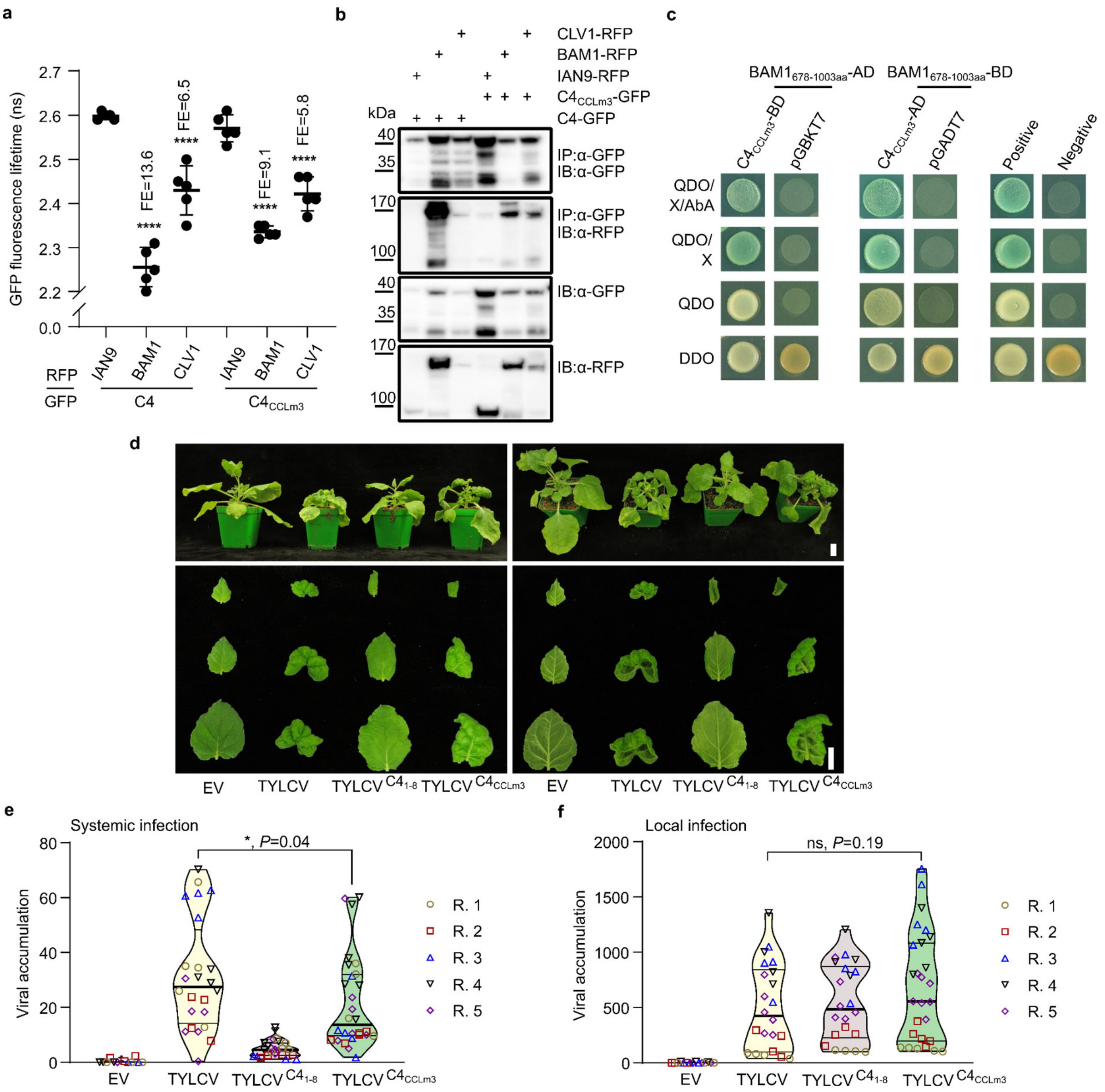


**Figure S17. $C4_{CCLm3}$ retains the interaction with plasma membrane-localized receptor kinases and induces milder symptoms than WT C4 in the context of the infection in *N. benthamiana*.**

a, b. Interaction between $C4/C4_{CCLm3}$ and the plasma membrane-localized receptor kinases BAM1 and CLV1 analyzed by FRET-FLIM (a) and Co-IP (b) upon transient co-expression in *N. benthamiana* leaves. Samples were taken at 2 days post-agroinfiltration (dpa). The interaction between C4 and BAM1 is used as a positive control. The interaction between C4 and IAN9 is used as a negative control. In (a), each dot in the graph represents the GFP fluorescence lifetime (ns, nanoseconds) obtained for one technical replicate consisting of one field; the thick line is the average value, and the error bars correspond to standard deviations. Significant differences between groups were determined by one-way ANOVA ($P<0.0001$, F=67.66, df=5), followed by multiple comparisons of means by applying Tukey test; asterisks represent statistically significant differences at: ****, $P<0.0001$; ns, not significant. FE, FRET efficiency. IP: immunopurification, IB: immunoblotting.

c. Interaction between the kinase domain (678-1003aa) of BAM1 and $C4_{CCLm3}$, tested by yeast two-hybrid (Y2H).

d. Representative phenotypes of *N. benthamiana* plants infected with TYLCV or TYLCV$^{C4_{CCLm3}}$ at 21 days post-inoculation (dpi). In upper panels, lateral and zenithal images of whole plants are presented; bottom panels show typical symptoms of infection in systemic leaves from three different plants. Scale bar, 2 cm.

e, f. Viral accumulation was analyzed in *N. benthamiana* plants in both systemic (e) and local (f) infections by qPCR, at 21 dpi (e) or 2 dpi (f). Violin plots show the aggregate results obtained in 5 independent trials, where each dot represents one biological replicate consisting of an individual plant, the thick lines represent median values, and thin lines the lower/upper quartiles. A minimum of 5 plants were analyzed per infection and trial, when possible. Differences between TYLCV and TYLCV$^{C4_{CCLm3}}$ were assessed by applying Mann-Whitney U test (*, $P$=0.042; ns, not significant). TYLCV$^{C4_{1\text{-}8}}$, a mutant virus carrying a premature stop codon mutation in the C4 sequence resulting in the translation of the first 8 aa only (Rosas-Diaz *et al*. 2018), was included in (d), (e) and (f) as reference; EV, Empty vector; R indicates each of the independent replicates.

The experiments in (a) and (b) were performed two times with similar results. The experiments in (c) were performed three times with similar results. One replicate is shown here.

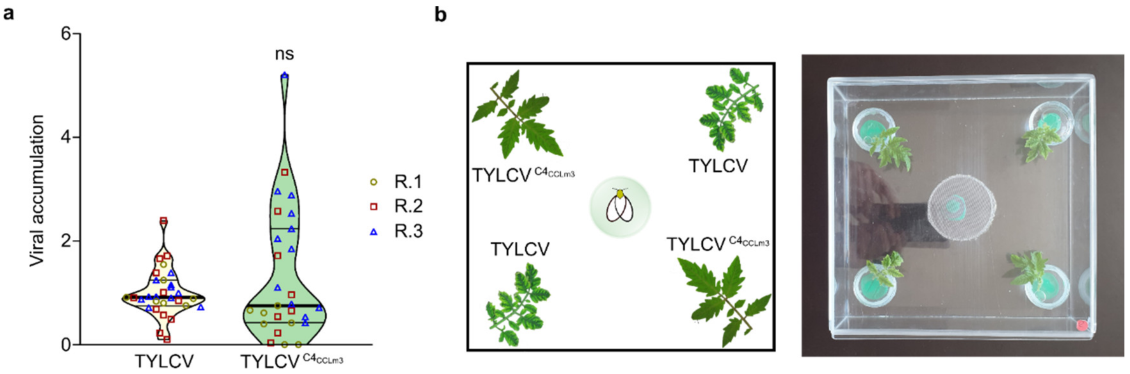


**Figure S18. Whitefly choice assay: viral accumulation and experimental design.**

a. Accumulation of TYLCV and its mutant form TYLCV$^{C4_{CCLm3}}$ in the tomato plants used for the dual choice assays with *Bemisia tabaci* (whitefly) in Fig. 4e. Systemic samples were collected at 19 days post-inoculation. The violin plot shows the aggregate results obtained in three independent replicates; each dot represents viral accumulation per individual plant, thick line the median, and thin lines the lower/upper quartiles. Statistically significant differences were assessed by applying Mann-Whitney U test; ns, not significant. R indicates each of the independent replicates.

b. Experimental design followed in dual choice assays with whiteflies and tomato plants infected with TYLCV or TYLCV$^{C4_{CCLm3}}$, as described in Ontiveros *et al*., 2022. TYLCV- and TYLCV$^{C4_{CCLm3}}$-infected leaflets were placed in the corners of a cage occupying alternate positions (2x2), surrounding the flight release platform situated in the middle, where an individual whitefly was placed and its preference recorded. Choice assays were repeated individually at least 60 times per experiment (60 individual whiteflies, i.e. 60 biological replicates).
